# High throughput quantitative tracking of *Plasmodium falciparum* clonal blood stage parasite growth and applications for antimalarial drug discovery

**DOI:** 10.1101/2025.11.30.691439

**Authors:** Zhe Cheng, Liyang Ru, Andrew Hoang, Mingyu Wang, Suryanarayana Ganjikunta, Steven P. Maher, Dennis E. Kyle

## Abstract

New systematic profiling of drug effects is in demand due to limitations in existing approaches to evaluate comprehensive drug effects and distinguish heterogeneity in mixed populations. One challenge is the underdeveloped methods for investigation of causal association between artemisinin induced dormancy and recrudescence in *Plasmodium falciparum*. We developed Quantitative Tracking After Chemotherapy Exposure (qTRACE) and an artificial intelligence (AI) mode to evaluate cytotoxic and cytostatic effects simultaneously at single parasite resolution in a high throughput platform. qTRACE is based upon the observation that individual parasites grow into colonies that can be quantified for numbers of viability and growth rate. Applying qTRACE, we revealed parasite-drug killing dynamics after artemisinin exposure, finding up to 50% of viable parasites arise directly from recovery of artemisinin induced dormancy. We further developed next generation qTRACE integrated with deep learning-based segmentation and analysis, thereby directly linking continuous, time-lapse phenotypes of label-free live *P. falciparum*. Our results confirmed the viability of dormant ring stages and that recovery rates differ between artemisinin-susceptible and -resistant *P. falciparum*.

## Main

Accurate evaluation of drug induced effects and parasite viability is essential for discovering new anti-malarial drugs and monitoring the success of currently used regimens. Previously, evaluation of susceptibility to anti-malarial drugs is measured by half-maximal inhibitory concentration (IC50) or various ring stage survival assays^1–5^. These assays are carried out in a bulk culture for short periods of time, after which susceptibility of drugs is measured by comparing the number of replicated parasites between treated and untreated condition. Many of these methods rely on taking aliquots from a large culture and only capture a single snapshot.

Drug perturbation assessments often lack insights into drug effects and are unable to differentiate growth inhibition and temporary arrest with killing. More critical limitations include the inability to evaluate long-term drug-parasite dynamics as the measurements are not derived from the same individual cells and cannot resolve heterogeneity in a mixed population. Too often the result is a misinterpretation of killing rate or that no viable parasites remain in the short-term assay. Conversely, previously published data suggest persistent survivors exist and that standard assays obscure true killing effects of drugs and hinder the development of next generation anti-malarial drugs^6^. Consequently, these methods create inconsistent correlations between drug susceptibility and genotypes, while ignoring valuable biological insights of individual diversity of parasites and drug effects in a continuous system.

Artemisinin combination therapy is the frontline treatment for *P. falciparum* infection. However, the emergence and spread of resistance to artemisinin poses a significant threat in malaria control^7^. Delayed clearance and recrudescence are frequently associated with artemisinin resistance clinically^8^. It has been widely reported that exposure to artemisinin drugs induces a phenotype known as artemisinin-induced dormancy and these dormant parasites can persist for many days before they either recover or die. Numerous studies suggest these dormant parasites play a role in infection recrudescence and treatment failures both clinically and in vitro^9–13^. Drug-induced cell cycle arrest has been widely observed in many human diseases, including malaria, and have posed a significant challenge in searching for improved therapies^6^. Thus, understanding the role of dormancy with artemisinin resistance and parasite recrudescence is critical for formulating effective counterstrategies. However, the causal relation of artemisinin induced dormancy to parasite survival and recrudescence remains poorly understood due to limitations in existing approaches. Some of the various assays attempt to compensate for delayed recovery and quantify dormancy recovery, but still face the same flaw that early survivors can easily disguise the existence of parasites that recover later in a mixed culture^2, 14^. An improved and more efficient evaluation of artemisinin effects and quantification of artemisinin induced dormancy are needed to accurately quantify drug resistance at the clonal level.

In this study, we developed Quantitative Tracking After Chemotherapy Exposure (qTRACE) to quantify viability and growth at single parasite resolution based on the concept that following an adequate dilution of parasitized erythrocytes, one parasite grows into a colony that can be visualized microscopically. qTRACE is a versatile method that simultaneously tracks survival, time-dependent recovery, and drug-induced growth effects. We demonstrated the robustness of qTRACE by revealing true long-term drug effects of dihydroartemisinin (DHA) on multiple *P. falciparum* isolates and cloned parasites with diverse backgrounds. We further clarify the underappreciated contribution of artemisinin induced dormancy in parasite recrudescence and reveal the undiscovered growth retardation effect of artemisinin drugs in selected parasite strains. Further development of qTRACE AI with deep learning-based cellular analysis effectively enhanced time-lapse biological connection, generalization, and accuracy through unrestricted observation of label-free live cell phenotypes. Applying qTRACE to drug screening, we found a diverse cytostatic and cytotoxic effect preference and time-dependent dynamics between drugs and parasites. qTRACE is strongly positioned for holistic investigation into single cell level drug-parasite dynamics while remaining amendable to high-throughput and logistically complex time course experiments.

## Results

### Quantification of longitudinal viability and growth of single parasites

We found that, when sufficiently diluted in a 384 well microtiter plate, one single parasite grows into a colony and can be continuously monitored over time. These initial observations led us to develop a high throughput and high content imaging assay. A previously developed GFP integrated parasite (NF54-GFP) was seeded into 384 well microtiter plates and imaged continuously using a 4x objective on a high content imaging system (HCI)^15^. In these assays, we observed continuous formation and growth of colonies over 22 days (Fig. 1A). Day 5 was the earliest time we could detect a colony, likely due to insufficient fluorescence intensity at low magnification at earlier time points. Due to the care taken to not disturb the plates during the time course, these colonies continued to grow at the same position in the well throughout the duration of the assay (22 days) (Fig. 1A). Meanwhile, we also observed parasite recovery and colony growth in ring stage *P. falciparum* exposed to 700nM DHA for 6h (Fig. 1A). Notably, some colonies appeared early, with others at later timepoints, which is consistent with the concept that artemisinin drugs induce ring stage dormancy, followed by delayed recovery of the parasites (Fig. 1A). Time-lapse imaging showed the differential onset of growth of individual parasites, with late recovering parasite colonies from dormancy detected in the same well alongside early recovering parasites (Supplementary Video 1). Comparison of GFP, Hoechst, and DIC imaging showed complete overlap of the identified colonies which confirms these colonies consist of parasites (Fig. 1B). We chose to use Hoechst-staining of DNA as the fluorescent marker as this route enables assessment of clinical isolates without integrated fluorescent markers. At increased magnification (60X and 100X), we show that a colony consists of hundreds-to-thousands of parasites, evidenced by imaging of individual parasites and foci of hemozoin (Fig. 1C). The growth of parasites results in reduced hemoglobin that appears as an opaque clearing in the red blood cell monolayer, consistent with previous study of plaque forming in *P. falciparum*^16^ (Fig. S1A). A z-stacked video of a colony imaged with a 100X objective shows that parasites are distributed horizontally and vertically throughout the packed red blood cell layer (Fig. S1B and Supplementary Video 2).

**Fig. 1.**
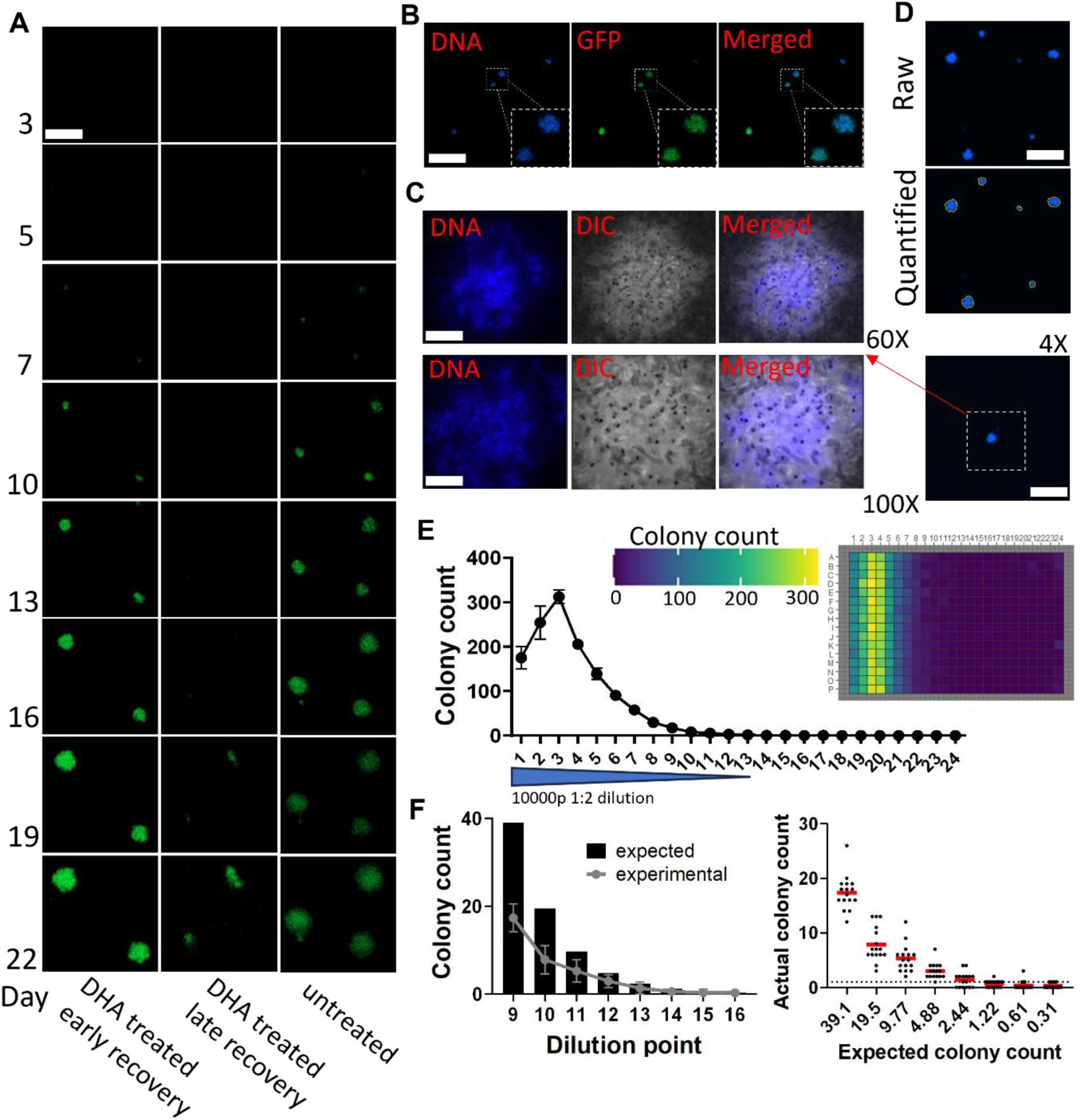
A single parasite grows into a colony. **(A)** Growth of the same individual NF54-GFP colonies imaged over time, with or without 6h DHA exposure (Scale bar = 150μm). **(B)** Hoechst stained and GFP fluorescent images of day 7 parasite colonies imaged with a 4X objective; white boxes show the same colonies imaged with a 20X objective (Scale bar = 100μm) **(C)** Higher magnification images of a day 7 parasite colony. The observed black dots are hemozoin. Top: imaged with a 60X objective (scale bar = 25μm), bottom: imaged with a 100X objective (scale bar =15μm). The 4X objective image scale bar = 100μm. **(D)** High-content image analysis was used to measure the number and the size of colonies of a fraction of a well (Full well is shown in Fig. S1C) (Scale bar = 100μm). **(E)** Average number of colonies in each column of each dilution point (left) and heatmap view of colony counts of the same serial dilution plate (right). Dilution points 1 and 2 could not be accurately quantified due to wells having too many colonies. **(F)** Comparison between experimental results and estimated numbers in low dilution points (left) and comparison of the number of colonies between quantified and expected number of positive wells (right). The estimated colony numbers are shown in Fig. S3A. All experiments were performed at least twice.

These phenotypes, therefore, can be used to quantify the growth of individual parasites (Fig. 1D and S1C). Long-term measurements at multiple time points allow for longitudinal measurements of parasite growth and identification of slow growing or late recovering parasites (Fig. S2). To validate the single parasite-to-colony concept and quantification reliability, we performed a serial dilution experiment to assess correlation between colony number and parasite input (Fig. S3A). Measurements of total DNA content showed a successful serial dilution of parasites (Fig. S3B). From the same experiment, quantification of colonies from wells seeded with ≤2500 cells correlated with the total DNA content of parasites; however, the first two dilution points resulted in overlapping colonies which lead to inaccurate quantification (Fig. 1E). In wells with lower numbers of parasites, the average quantified colonies in each well were within one dilution of the expected number of colonies (Fig. 1F). Taken together, these results demonstrate that one colony comes from replication of one single parasite and quantification of colonies can be used to track the original number of viable parasites in each well.

### qTRACE tracks single parasite survival, time-dependent recovery and growth

Initially, we developed qTRACE to improve our understanding of survival and recovery from artemisinin exposure and artemisinin-induced ring stage dormancy phenotypes (Fig. 2A). To ensure sufficient identification of late recovery of dormant parasites, we designed the assay to last 22 days in duration, with multiple plates prepared for multiple timepoints throughout (Fig. 2A). To minimize liquid movement in the blood layer and to preserve colony integrity, we designed a robotic automation protocol to perform slow liquid manipulation with essentially no disruption to the blood layer (Figure S4A). At every timepoint, one plate was fixed and stained for HCI while the remaining plates underwent media change using the designed no-disruption protocol (Fig. 2A).

**Fig. 2.**
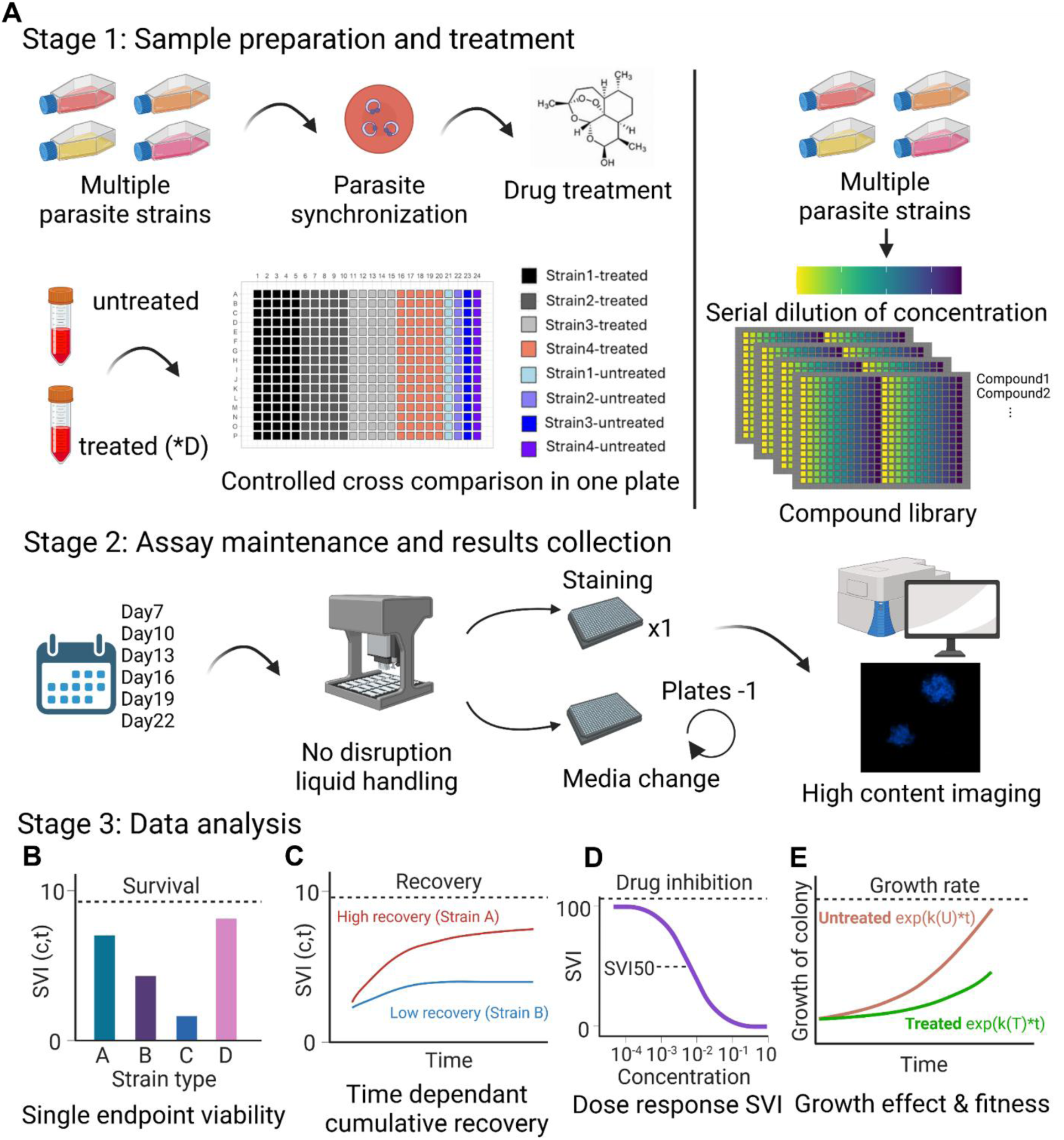
qTRACE simultaneously reports viability, time-dependent recovery, and growth effects at single parasite resolution. **(A)** Schematics of the qTRACE process. In stage 1, parasite strains are prepared and treated with drugs. Treated and untreated samples are prepared based on the dilution factor D, then plated on the same plate. Left side showing representative DHA exposure workflow and right side showing representative drug testing workflow. In stage 2, plates undergo automated media change, and one plate is taken for HCI at each designated timepoint. In Stage 3, data are analyzed as described in Fig. 2B-E. **(B)** Single endpoint SVI assessments to evaluate percentage viability at concentration c and time t. **(C)** Assessments of SVI at multiple time points to evaluate time-dependent recovery or drug effects. **(D)** SVI assessments in a dose response. SVI50 is the concentration of a drug that estimates 50% of parasites were viable. **(E)** Assessments of exponential growth of colonies under different conditions or between different strains to evaluate growth effects and fitness. Growth Rate (GR) is then derived using variable K between two conditions.

We developed a single-parasite viability index (SVI) to compute the viability from drug treatment based on quantification of colonies. We can determine the viability of single parasites at time t for strain s:

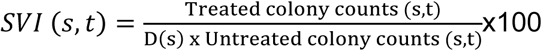

SVI is the ratio of the number of colonies between treated and untreated conditions normalized to single parasite division. D(s) is the determined dilution factor between treated sample and untreated sample for strain s. The introduction of factor D enables greater tolerance of experimental variability caused by cell density (Fig. S4B). Additionally, the factor D can be adjusted to compensate for different levels of drug sensitivity of different genotypes. SVI showed minimal variation under different D factors because the overall percentage of viability remained unchanged (Fig. S4C).

The length of qTRACE experiments is flexible and a single timepoint qTRACE SVI assessment is sufficient for evaluating drug killing and parasite survival, which requires no media change and can be applied with manual manipulation (e.g., multi-channel pipettes) (Fig. 2B). In this study, we used day 7 as the timepoint to assess initial survival from artemisinin exposure. By adding time points to generate time-dependent SVI, we observed cumulative recovery overtime at single parasite resolution (Fig. 2C). This made it possible to identify and quantify parasites that recovered late. In addition to evaluation of multiple parasite strains treated with a single inhibitor, qTRACE also is compatible with high-throughput drug screening (Fig. 2A). In a dose response experiment, we assessed drug killing effects at different concentrations (Fig. 2D).

Through quantification of the colony size over time, we captured the exponential growth of viable parasites (Fig. 2E). These data are useful to evaluate effects of drug on parasite growth, making cytostatic effects measurements a convenient secondary outcome from qTRACE. Moreover, quantification of the area of colonies alongside viability enables identification of late recovered dormant parasites and assessment of growth behavior of those recovered parasites. To quantitatively determine growth difference between two samples, we adapted the growth rate inhibition metrics (GR) from a previous study^17^:

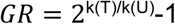

In our application, k(T) is the growth rate of drug-treated parasites and k(U) is the growth rate of untreated controls. Similarly, the GR value can also be used to quantitatively assess fitness difference between strains, where k(T) was the growth rate of strain 1 and k(U) was the growth rate of strain 2. The GR value ranges from -1 to 1, with 1 suggesting no impact on growth, 0 meaning complete growth inhibition and negative value indicating cytotoxic effects. Using GR and SVI together we differentiated between cytotoxic and cytostatic effects of a drug.

### qTRACE reveals time-dependent viability of single parasites after DHA exposure and diverse mechanisms of artemisinin induced dormancy

Mutations in the *P. falciparum* K13 propeller domain are known to facilitate resistance to artemisinin drugs^18^. To study how K13 mutations affect the viability of parasites, we compared Dd2 with its K13 mutants Dd2-K13-C580Y and Dd2-K13-R539T using qTRACE^19^. Rings at 0-3hpi were exposed to 700nM DNA for 6 hours before plating (Fig. 2A). As expected, we captured the number and the size of growing colonies from all three strains and observed significantly fewer colonies from DHA treated Dd2 compared to its isogenic K13 mutants (Fig. 3A). Overall, both DHA treated and untreated colonies kept growing throughout the assay duration (Fig. S5A). Interestingly, a wide distribution of colony sizes was observed even in synchronized untreated controls, presumably due to genuine growth variability as previously noted^20^ (Fig. S5A). qTRACE single timepoint assessment of SVI on day 7 revealed differences in susceptibility to DHA among three strains (Fig. 3B). All three strains continued to show increased viable parasites over time until reaching a plateau, indicating that the initial survivors only represent a fraction of the total parasites that eventually recover following exposure to DHA (Fig. 3C). These results demonstrate that both Dd2 and its K13 mutants exhibit dormancy phenotype for survival and that mutated K13 facilitated resistance is manifested on both initial survivors and recovery from dormancy mechanisms. Dd2-K13-R539T exhibited over 100-fold more viable parasites after exposure to DHA, whereas Dd2-K13-C580Y showed a comparatively lower difference compared to the parental Dd2 (Fig. 3D). This is consistent with previous reports showing that the R539T mutation confers a higher level of resistance than C580Y^18, 21^. The fold change remained stable for all three strains, indicating that although higher number of parasites recovered in K13 mutans, the proportion of late recovery remained similar across three strains.

**Fig. 3.**
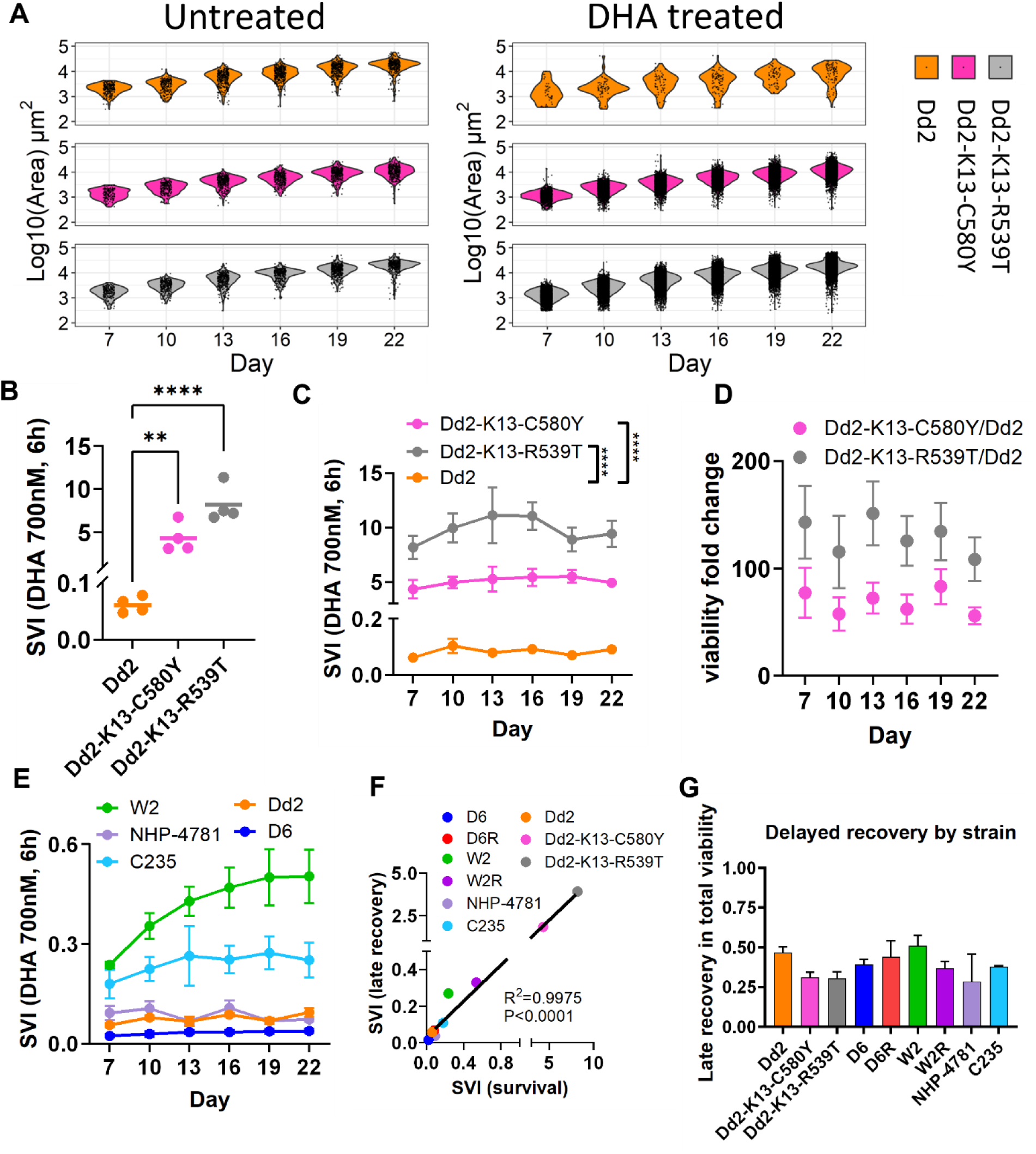
qTRACE reveals differential susceptibility and recovery from DHA exposure across various parasite lines. **(A)** Individual colonies and the sizes of each colony under untreated condition (left) and DHA condition (right) for every timepoint recorded from Dd2 and Dd2 K13 mutants. Each black dot represents one colony. From two biological replicates. **(B)** SVI assessment of Dd2 and Dd2 K13 mutants on day 7 quantifying initial survival from DHA exposure, n=80 technical replicates and n=4 biological replicates. P-value calculated from one-way Anova. **(C)** Time-dependent SVI of Dd2 and Dd2 K13 mutants at continuous timepoints, day 19 and 22 for Dd2-K13-R539T were not accurately quantified due to colony overgrowth. n=80 technical replicates and n=4 biological replicates. P-value calculated from two-way Anova. **(D)** The fold change in SVI between Dd2 and Dd2-K13-C580Y, as well as between Dd2 and Dd2-K13-R539T, for all timepoints. **(E)** Time dependent recovery SVI of five artemisinin sensitive strains at continuous timepoints. n=80 technical replicates and n=3 biological replicates. **(F)** Comparison of late recovery SVI (Max SVI-survival SVI) and survival SVI of all nine parasite lines with linear regression. P-value and R^2^ value calculated using Pearson’s correlation. **(G)** Proportion of late recovery in total viability over entire 22 days across nine parasite lines (error bar=SEM). Additional data in Supplementary Table 1.

To investigate if the delayed recovery from DHA is a common strategy of survival, we tested various other laboratory-adapted *P. falciparum* lines and clinical isolates. We tested two parasite lines and their progenies that were previously selected for artemisinin resistance (D6, D6R, W2 and W2R)^9^. Both W2/W2R and D6/D6R pairs showed different levels of resistance to DHA exposure and continued to have parasites recover over time (Fig. S5B and S5C). The decreased fold change of viability between W2 and W2R shows that W2 has more viable parasites coming from delayed recovery than W2R, presumably because W2R adapted stronger DHA tolerance during resistance selection (Fig. S5B). The viability difference between D6 and D6R was greater than that between W2 and W2R, presumably because D6R went through a longer selection period than W2R (Fig. S5C)^9^. Overall, the in vitro drug selected parasites exhibited lower levels of resistance compared to parasites with mutated K13, yet recovery from artemisinin-induced dormancy was observed in both genotypes (Fig. 3C, S5B and S5C). We next tested NHP-4781 which has E252Q K13 mutation that is located outside β-propeller domains^22^. NHP-4781 showed a comparative level of susceptibility to DHA exposure compared to Dd2, indicating that mutation outside propeller domain on K13 does not incur resistance to artemisinin, which is consistent with previous findings^21^ (Fig. 3E and Fig. S5D).

TM91-C235 is a multi-drug-resistant parasite derived from patient with multiple drug treatment failures^23^; it showed moderate resistance compared to D6 and Dd2, but not to W2 (Fig. 3E and Fig. S5D). The ability to have more continuous dormancy recovery of W2 compared to other parasite strains is consistent with previous study which showed higher and longer recovery of W2 after DHA exposure (Fig. 3E)^14^.

The nine *P. falciparum* lines tested showed different levels of delayed recovery from DHA (Fig. S5E). Notably, the initial survival measured on day 7 from DHA strongly correlates with late recovery, indicating that, despite the differences of susceptibility to DHA among these parasites, the survival is equally exhibited by the initially survived parasites and parasites recovered from dormancy (Fig. 3F). For many of the artemisinin sensitive parasite lines, delayed recovery parasites accounted for about half of the total parasites that eventually survived DHA exposure (Fig. 3G). A decreased proportion of late recovered parasites in both K13 mutants and in vitro selected lines suggests that an increased level of resistance reduces the reliance on dormancy as a mechanism to survive (Fig. 3G). The large portion of delayed recovered parasites is often overlooked in many studies that use standard 72-hour assays to estimate parasite killing but now can be more accessibly evaluated using qTRACE.

### qTRACE assesses growth inhibition and fitness

Through measurements of the growing colonies, qTRACE allowed us to quantify growth inhibition imposed by drugs and fitness differences between parasite lines. From the same DHA exposure experiments described above, we analyzed the growth and distribution of colonies of DHA treated Dd2 and its K13 mutants. We found the growth of the two K13 mutants mimics their untreated control, while the growth of Dd2 was severely affected by DHA, as evidenced by the dispersed distribution (Fig. 4A). Comparison of growth curve and GR value showed Dd2 suffered a reduced growth rate after DHA exposure, whereas K13 mutants showed overlapped growth with their control (Fig. 4B and 4C). These results suggest that Dd2 K13 mutant parasites can quickly adapt to normal growth once recovered from DHA, but the growth of many parental Dd2 parasites become impaired by DHA. Many previous studies showed that adaptation to drug resistance sometimes induces fitness costs in the parasites^10, 24^. Through qTRACE, we can detect fitness disadvantage in the Dd2-K13-C580Y, but not in the Dd2-K13-R539T, compared to parental Dd2; which is consistent with previous reports^21^ (Fig. 4D). In the case of in vitro selected lines, W2R showed a fitness disadvantage compared to W2, while D6R showed a more minor disadvantage compared to D6 (Fig. S6A), as noted previously^10^. DHA exposure resulted in reduced growth and lowered GR for W2 and D6, yet W2R and D6R phenocopied the growth of untreated conditions (Fig. S6B and S6C). C235 showed a slight impact on growth rate from DHA exposure (Fig. S6D). NHP-4781 also exhibited retarded growth following DHA exposure, as well as poor adaptation to culture as seen by the slow growth in the untreated control (Fig. S6E). This is supported by the observed gametocyte formation even during routine culture. Total viability from artemisinin exposure showed a moderate correlation with GR, indicating that increased resistance to artemisinin led to decreasing growth impact on the viable parasites, but was also parasite background dependent (Fig. 4E). Together, these results demonstrate that resistance to artemisinin is also reflected in the ability to quickly resume growth upon recovery.

**Fig. 4.**
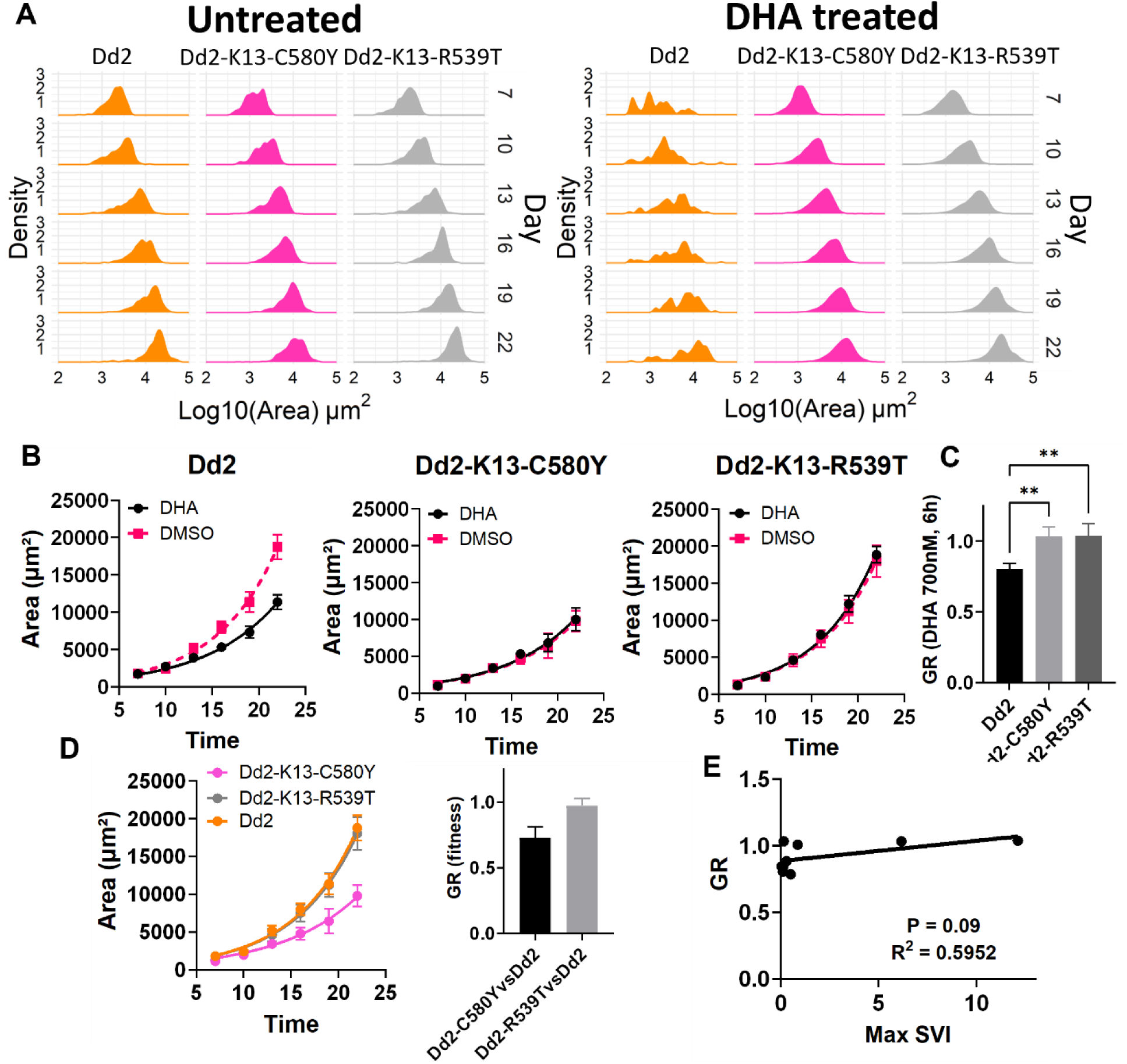
Growth impacts induced by DHA and fitness costs to artemisinin resistance. **(A)** Density distribution of colony sizes in Dd2 and Dd2 K13 mutants at continuous timepoints in DHA and DMSO conditions. From two biological replicates. **(B)** Fitted growth curves of averaged colony sizes at all timepoints for Dd2 (left), Dd2-K13-C580Y (middle), and Dd2-K13-R539T (right) in DHA condition and DMSO condition. n=80 technical replicates and n=4 biological replicates. **(C)** GR values of Dd2 and Dd2 K13 mutants comparing DHA condition to DMSO condition for each strain. P-value calculated from one-way Anova. **(D)** Fitted growth curves of averaged colony sizes at all timepoints for Dd2 and Dd2 K13 mutants without any treatment (left) and GR values between parasite lines (right). n=80 technical replicates and n=4 biological replicates. **(E)** Correlation comparison between Max SVI and GR for all nine parasite lines tested with linear regression. P-value and R^2^ value calculated using Pearson’s correlation. Additional data in Supplementary Table 2.

### Application of qTRACE in drug testing

The ability to assess phenotypes of single parasites is a valuable tool to improve accuracy of drug susceptibility testing (Fig. 2A). We applied qTRACE to drug screening and tested 15 common anti-malarial drugs on Dd2 parasites (Fig. S7A and Supplementary Table 3). Consistent with previous reports of Dd2 background, qTRACE also demonstrated elevated resistance to sulfadoxine and pyrimethamine in Dd2^25^. The SVI50 of most compounds tested showed comparable results to the standard IC50 measurements in a bulk culture, indicating strong reproducibility of qTRACE to serve as a quantitative alternative to conventional bulk methods (Fig. S7B and Supplementary Table 3). qTRACE also demonstrated extraordinary efficiency and improvements of throughput due to very low parasite input requirements (less than 50 parasites per well).

### qTRACE AI powered by deep learning enables continuous label-free live cell analysis

From the development of qTRACE with DNA dyes, we noted one of the major limitations is the lack of continuity between every timepoint in a continuous experiment when not using self-reporting parasite isolates (e.g., GFP) (Fig. 2A). To enable continuous observation of the same samples and preserve clone identities, we incorporated a deep learning-based semantic segmentation method into qTRACE. The next generation qTRACE AI directly quantified objects on images acquired through transmitted light microscopy without the need to fix, stain, or genetically modify the parasites. A total of 5760 distribution-calibrated images spanning diverse experimental applications were used to train convolutional neural networks (CNN) (Fig. 5A).

**Fig. 5.**
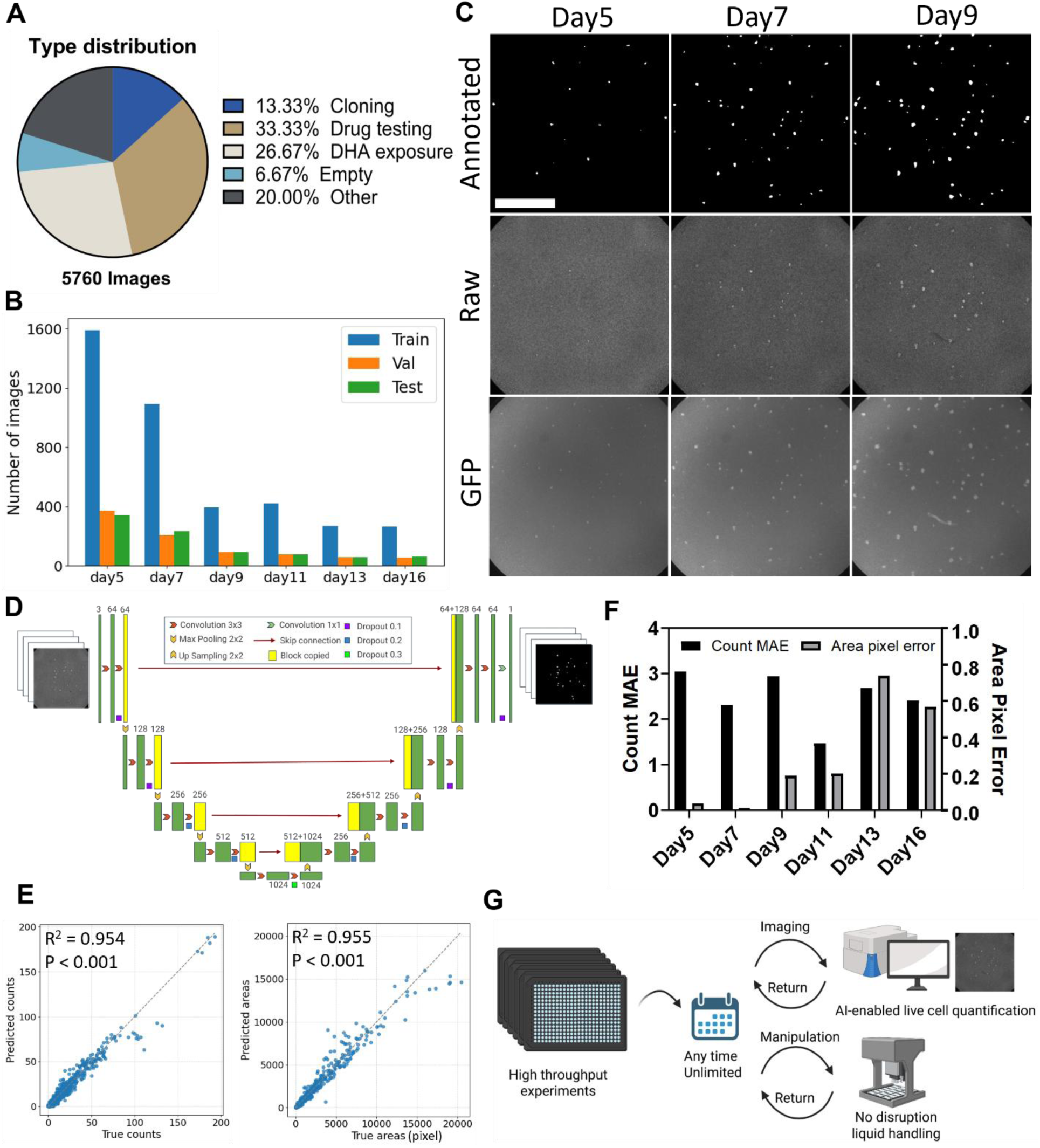
Deep learning enables continuous label-free live cell analysis. **(A)** Distribution of training datasets by types of experiments. **(B)** Distribution of training datasets by time points and training, validation, and test. **(C)** Representative raw transmitted light images and human annotated mask images for a continuous experiment. GFP was used as annotation reference to confirm authenticity of colonies on transmitted light images. Scale bar 1000μm. **(D)** Schematics of model architecture and qTRACE AI application. The architecture consists of a contracting path (left side) and an expanding path (right side) with skip connections (red arrows) that concatenate feature maps between corresponding layers. Numbers indicate the spatial dimensions and number of feature channels at each layer. **(E)** Evaluation of model prediction accuracy compared to human annotated results for count (left) and area (right). P-value and R^2^ value calculated using Pearson’s correlation. **(F)** Count mean absolute error (MAE) and area pixel error percentage of the trained model by timepoint. **(G)** Schematics showing qTRACE AI workflow. Flexible continuous imaging and manipulation of the same sample over time.

These images covered different time points and sizes of colonies, but with a stronger focus on early times to compensate for increased difficulty of learning to differentiate small colonies (Fig. 5B and S8A). Parasite colonies of every transmitted light image were annotated to polygons by trained human professionals using ground truth GFP images as reference (Fig. 5C).

We trained on CNN model U-Net architecture because of its established capability of biomedical level semantic segmentation (Fig. 5D and S8B). Overall, qTRACE AI achieved comparable human level accuracy, demonstrated by the strong correlation between annotated results and model predicted results (Fig. 5E). Mean absolute error (MAE) of count quantification showed an average of 2.63 and is relatively stable over time, while area pixel error increased slightly in later timepoints (Fig. 5F and Supplementary Table 4). Most of the test images showed identical quantification between annotated results and predicted results across various conditions (Fig. S9A). However, the true accuracy of qTRACE AI is underestimated here because we noticed most of the disparities between annotated results and predicted results were due to human errors in annotations. In almost all instances, our trained CNN model was able to correct human errors, including filtering debris and false positives, and to identify true positive colonies that were originally omitted by human observations (Fig. S9B and S9C).

Additionally, the label-free image analysis of qTRACE AI removed the most common challenge of dealing with fluorescent noise during imaging as we routinely saw images that would have been unsuitable for analysis due to high fluorescent background (Fig. S9D). Since qTRACE AI does not have any benchmarking available, we decided to train a second model using Vision Transformers (ViT) to allow for a performance comparison (Supplementary Table 4). Overall, U-Net architecture showed superior performance compared to ViT for both count and area quantification tasks (Supplementary Table 4). The performance differences between architectures may be attributed to the data-efficiency characteristics of each model. Thus far, the capability of qTRACE AI to track live single parasites greatly enhances accuracy and flexibility over DNA stain qTRACE protocol, making it more suitable for diverse applications, including demanding long-term studies that may yield new insights into drug effects (Fig. 5G).

### qTRACE reveals new insights into drug activities and effects

We screened a Medicines for Malaria Venture (MMV) antimalarial reference library to derive both SVI50 and GR50 using qTRACE AI label-free imaging analysis (Fig. 6A). A correlation study of qTRACE AI and qTRACE side by side showed excellent reproducibility between the two methods for viability and growth assessments (Fig. 6B). Most of the tested drugs showed similar dose response curves between qTRACE AI and qTRACE (Fig. S10A).

**Fig. 6.**
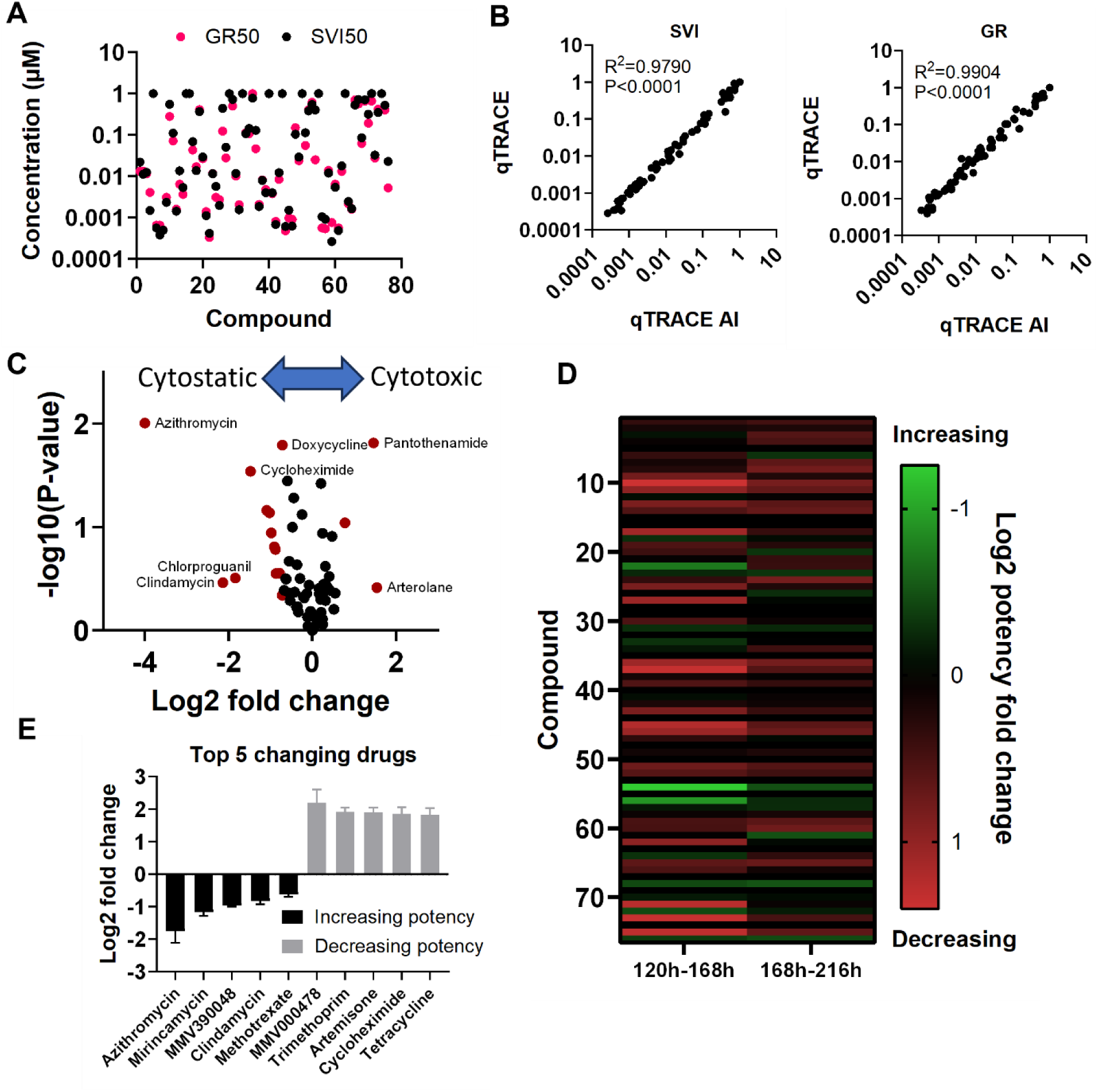
qTRACE reveals new insights into drug activities and effects. **(A)** SVI50 and GR50 for drugs tested from MMV antimalarial reference library. n=2 biological replicates. **(B)** Comparison of screening results between qTRACE AI and qTRACE for SVI (left) and GR (right). P-value and R^2^ value calculated using Pearson’s correlation. **(C)** Cytotoxic and cytostatic effect evaluation of drugs from MMV antimalarial reference library. Shifting in directions represents preference of either cytostatic effect (left) or cytotoxic effect (right). **(D)** Effect of altering time on SVI50 shift for drugs from MMV antimalarial reference library at 48h interval. n=2 biological replicates. **(E)** The five drugs showing the highest potency shift, for each direction, over entire measured time course are shown. Error bar = SEM.

These results confirmed the feasibility of qTRACE AI to obtain validated results from more typical experimental applications. Through the evaluation of SVI50 and GR50, we identified diverse drug induced cytotoxic and cytostatic effects from tested compounds. (Fig. 6C and Supplementary Data 1). Azithromycin showed strong growth inhibition evidenced by the much lower GR50 and the inhibition of growth at lower concentration as seen in the GR curve, while pantothenamide showed clear cytotoxic effect (Fig. 6C and S10B). Understanding time-dependent drug effects is crucial for dosing and pharmacokinetic studies. To take advantage of the unlimited continuous live cell imaging of qTRACE AI, we performed a time-lapse experiment to assess the time-dependent potency of compounds from the same cells over time. We observed all three types of time-dependent activities, either increasing, decreasing or stable over time (Fig. 6D). We detected 23 drugs that gradually lost potency, while only 3 drugs gained extra potency with at least 2-fold change over time (Fig. 6D and Supplementary Data 1).

Azithromycin showed the greatest potency increase in the library, as well as mirincamycin, another known slow acting drug^26, 27^ (Fig. 6E). KAE609, KAF156 and some other drugs showed strong stability over time, while many of the drugs lose their effect over time (Fig. 6E and Supplementary Data 1). From these results, we conclude that qTRACE is capable of being used as a significant enhancement over conventional drug testing to reveal more insightful drug effects at single parasite resolution.

## Discussion

The development of qTRACE was originally driven by the ambition to elucidate the confounders in artemisinin induced dormancy and to quantify viable parasites that result in recrudescence. Numerous studies have suggested the association between artemisinin resistance and artemisinin induced dormancy, yet there have been limitations in quantifying viable parasites that recover later than the first recrudescent brood^9, 10, 12, 14, 28^. qTRACE overcomes those limitations by tracking the viability and growth of individual parasites, thereby effectively distinguishing initial survivors and late recovered parasites in an automated high throughput manner. Our results confirmed that recovery from artemisinin-induced dormancy plays a significant role in overall parasite survival at the population level, with nearly half of the viable parasites arising from dormancy recovery. Given that day 7 was the first point of measurement, the proportion of delayed recovery is likely underestimated. In this study, we show that the ability to survive artemisinin drugs in K13 mutants are not only exhibited by the large number of initial survivals, but also in the delayed recovery from dormancy (Fig. 3C, 3F and 3G). In fact, the number of parasites recovered from dormancy could be underestimated due to limited space in the well for higher resistant lines (both Dd2 K13 mutants) in later timepoints (Fig. 3C). The ability to utilize dormancy as a survival mechanism in both sensitive and resistant parasites suggest dormancy is an innate survival mechanism and likely independent of genomic resistance mechanisms. Resistance mechanisms improve initial adaptation and stress response to artemisinin drugs through each of their pathways, allowing more parasites to survive as well as more parasites to utilize dormancy. On the other hand, more sensitive parasites have overall lower initial survival and lower dormancy recovery rates due to poor initial response to artemisinin. Previous study suggests that dormant parasites are resilient to a wide spectrum of anti-malarial drugs, indicating the reduced efficacy of ACT partner drugs to remove dormant parasites^12^. Given the large portion of late recovery after DHA exposure, it would be reasonable to attribute the clinically observed recrudescence to the recovery of dormant parasites. In fact, parasites going into dormancy may be a favored survival path during ACT treatment because of the reduced susceptibility to drugs and the half-lives of the partner drug.

Investigation of cytostatic drug effects and fitness costs associated with increased resistance constitute a large study domain in drug discovery and drug resistance. The cytostatic effects of many anti-malarial drugs remain poorly studied due to lack of approaches to properly evaluate them in a large scale. Our ability to track individual parasites with qTRACE made it possible to evaluate cytostatic effects in a high throughput and technically straightforward manner. Across nine parasite lines, we observed varying cytostatic effects induced by artemisinin, influenced by genetic backgrounds and a moderate correlation of decreased growth inhibition to increased resistance.

Altogether, our results demonstrate that resistance to artemisinin is a multifaceted phenotype. When resistance increases, the proportion of initial survival and recovery from dormancy equally increases, meanwhile the impact on growth gradually decreases until there is no more growth impact, but in a parasite background dependent manner. The dormancy effects of artemisinin drugs cannot be underestimated and is likely the ultimate culprit behind frequently reported recrudescence. It is imperative that future ACT development and artemisinin resistance studies incorporate dormancy recovery as a standard evaluation.

Previous studies on fitness typically include laborious processes and error prone techniques, such as large amount of microscopy counting or combining two different parasites into one culture^10, 24, 29^. In this study, we showed that fitness assessment is a convenient secondary outcome when conducting qTRACE for other purposes, such as testing inhibitor effects. With the emergence of qTRACE, fitness studies will no longer need to be a complicated independent experiment. The growth assessment capability of qTRACE will be useful to a wide range of research, including but not limited to evaluating growth competition and growth inhibition, such as one imposed by RNAi or nutrition starvation.

Integration of deep learning-based analysis into qTRACE AI effectively overcame the limitation of having disconnected results at every timepoint. The U-Net architecture CNN model is well suited for qTRACE, evidenced by strong learning of the pattern with limited datasets. We found that qTRACE AI is superior to DNA stain qTRACE for its excellent generalization, accuracy, and cellular information linkage. qTRACE AI uses a single pipeline for all the experimental conditions, while in many cases different sets of rules are needed for DNA stain qTRACE depending on the timepoints or applications. Debris and background signals are among the most difficult noises to remove in rule-based image analysis because most of the time they fall into similar characteristic regions to the true objects. qTRACE AI demonstrated strong ability to handle those difficult situations (Fig. S9B, S9C and S9D). Lastly, label-free continuous live cell observation ensured the most authentic connection of cell behavior over time, and significantly improved our ability to identify unique signatures for each individual parasite. qTRACE AI and original DNA stain qTRACE will serve as great complement for each other to improve versatility and accessibility.

When applied to drug screening, qTRACE demonstrated its ability to serve as a direct replacement to currently used dose response assays. SVI50 and GR50 derived from qTRACE are robust alternatives to IC50 as the measurements were derived on a per parasite basis. For routine drug testing, a single endpoint at 120h is sufficient to identify different effects and does not require any robotic automation. Taking advantage of deep learning-based continuous label-free imaging, we screened an anti-malarial drug reference library and revealed new insights into both cytotoxic and cytostatic effects imposed by drugs, which will improve understanding of the mode of action of a drug and its future development. Through continuous measurements, qTRACE outputs time-dependent drug potency shifts, which can serve as foundation for drug pharmacodynamics and drug combination studies. Equally important to accurate drug effect evaluation is improved efficiency; qTRACE increases drug screening throughput by up to 667-fold. Typical dose response assay uses >1% parasitemia culture (∼2000 parasites/µl in 2% hematocrit), while qTRACE only needs less than 3 parasites/µl.

qTRACE is based on the biological phenomenon that a single parasite grows into a colony. The concept of colonies and plaque assays have been widely used in various biological research. In malaria research, similar concepts have been previously proposed, yet there has been little application and requires visual inspection of plaques^16^. Integrated with HCI, automation and deep learning, we developed a versatile application of this biological phenomenon and removed many of the previously suggested limitations, such as long waiting time and hematocrit limits. The flexibility to change media and supplement fresh blood is crucial for any long duration experiments. Here we have demonstrated that currently available compact automation stations are sufficient to achieve media change without disrupting colony integration, though it is not required for many of the applications shown in the report. Potential applications of this concept are far beyond what has been demonstrated in this study. Because of the ability to identify the number of colonies in each well, it becomes possible to map genomic and transcriptomic signatures associated with differential conditions at monoclonal levels. More advanced utilization could improve the selection of drug resistance by picking parasites that recover early or recover late, depending on if the purpose was to select for initial survivors or study latency mechanism(s).

In summary, qTRACE will serve as an all-around method that can be used for a wide range of scenarios, effectively overcoming previous limitations to achieve single parasite level insights. It would be appropriate to use qTRACE as a routine assessment of the recrudescence risks of developing compounds and current treatment combinations.

## Methods

### Parasite culture

The parasites used in this study include Dd2, Dd2-K13-C580Y and Dd2-K13-R539T (genetically modified Dd2 strains containing mutation on K13 PF3D7_1343700); D6 and D6R (an in vitro selected artemisinin resistant progeny of D6); W2 and W2R (an in vitro selected artemisinin resistant progeny of W2); C235; NHP-4781; and a genetically modified NF54-GFP (inserted GFP in cg6 locus)^9, 15, 19, 23^. For routine culture, parasites were grown at 37°C at 2% hematocrit in RPMI 1640 media (Gibco) supplemented with 25mM HEPES and 24mM NaHCO_3_, and with 10% heat-inactivated human A+ plasma and 10 µg/ml gentamycin. Cultures were maintained in a gas chamber with 5% O_2_, 5% CO_2_, and 90% N_2_. Giemsa slides were routinely made to assess parasite growth and stages. The growth media was changed every two days, and fresh blood was added when needed. Cultures were tested and found negative for mycoplasma regularly during this study.

### Ring Stage 0-3hpi Synchronization

The same 2-step synchronization protocol was performed to achieve 0-3h post invasion (hpi) for all strains used in this study. First, late stages were selected by laying on top of 65% percoll gradient and followed by centrifugation at 1500RCF^30^. Three hours later, freshly invaded rings were selected using 5% D-sorbitol ^31^. After each synchronization step, parasites were washed with RPMI 1640 media. Synchronized 0-3hpi parasites were allowed to reinvade before being used for experiments.

### Dihydroartemisinin (DHA) exposure

The same DHA treatment was used for all DHA exposure experiments in this study. Tightly synchronized 0-3hpi parasites were adjusted to 1% parasitemia and 2% hematocrit and treated with 700nM DHA. Six hours later, DHA was removed by centrifugation and washing with RPMI 1640 media before being replaced with culture media.

### GFP live cell time lapse imaging and fluorescent imaging

For GFP continuous imaging, 0-3hpi NF54-GFP parasites were seeded into 384 well plates (Greiner Bio-one) with an estimated number of 10 parasites per well. The same plate was taken for imaging on day 3, 5, 7, 10, 13, 16, 19, 22 using Lionheart (Agilent). Media was changed at day 7, 10, 13, 16, 19 by removing and replacing 20µl of media using NXP workstation (Beckman). Another experiment with 6h DHA exposure was performed in a similar way except the initial plated parasites were 1000 parasites per well. To image parasite using both Hoechst staining of DNA and GFP expression, 40ul of 5ug/ml Hoechst was added into the plate and maintained for 30 minutes before imaging. High magnification images were obtained using the 100x objective on a DeltaVision II (GE Healthcare) with deconvolution of a 22-plane Z-stack using Software (GE healthcare).

### qTRACE

#### Metrics

Colony counts and colony areas in the presence of drug are normalized to untreated controls in the same plate under the same growth conditions. Technical replicates were averaged to yield an average colony counts or colony area. For single candidate testing, we typically collect from 48-176 technical replicates. For dose response, two technical replicates per plate were collected.

Single-parasite viability index (SVI) was calculated to evaluate the viability of parasites and the cytotoxic effects of the testing drugs. SVI of parasite strain (s) at timepoint (t) was calculated as:

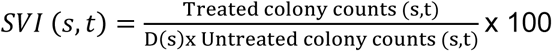

The average number of colonies for each parasite strain growing at each condition at every timepoint was calculated and normalized to the average number of colonies from the control. (D) represents the dilution factor between treated and untreated in a multi-strain testing design. SVI is a percentage value and ranges from 0 to 100, 0 meaning no viable parasite exists and 100 means no effect on parasite viability from the treatment condition. In recovery experiments, Survival SVI is defined as the SVI form the first point of measurement. The Max SVI measurement after the first timepoint is defined as the maximum possible viability or max SVI. Maximum delayed recovery is calculated as Max SVI – Survival SVI. When calculating SVI for a range of concentrations, SVI50 is the concentration that confers half maximal drug effect on parasite viability.

Growth rate inhibition (GR) was adapted from a previous study to evaluate the growth of every viable parasite^17^. The average area of colonies for each parasite strain growing at each condition at every timepoint was calculated and normalized to the average area of colonies from control. GR value ranges from -1 to 1, with 1 meaning no impact on growth, 0 meaning complete growth inhibition, and -1 meaning highest cytotoxicity. When calculating GR for a range of concentrations, GR50 is the concentration when GR = 0.5, which refers to half maximal growth inhibition.

For continuous measurements from multiple timepoints, an exponential growth curve was fitted for each parasite strain and each condition. GR was then calculated:

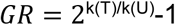

The growth rate (k) of each fitted curve was extracted, with k(T) representing the growth rate of drug treated condition and k(U) representing the growth rate of untreated controls. Alternatively, when testing fitness or growth competition between two strains, k(T) became the growth rate of strain 1 and k(U) became the growth rate of strain 2 or reference strain.

When measuring qTRACE as a single timepoint experiment, GR was calculated as:

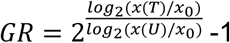

In this equation, x(T) represents the average area in drug treated conditions and x(U) is the average area in untreated control. X_0_ is the starting area; this could be the measurement before starting the experiment or be fitted as an estimated area of a single parasite because colonies originate from single replicating parasites.

### Single candidate qTRACE

This method was designed to test the response of multiple parasite lines to a single testing candidate of interest. In this study, DHA was used as an example. Synchronized 0-3hpi parasites were treated with DHA for 6h using the procedure described above. An estimated number of 10-50 parasites per well were seeded for untreated control for each parasite line. Depending on the designed D factor for each parasite line, D*number of parasites in untreated controls for each treated samples were seeded into the same plates with the untreated controls based on similar design shown in Fig. 2A. Based on different timeline design, at every timepoint, one plate was taken for staining and thereafter imaging quantification; the rest of the plates were taken for media change using designed no disruption protocol on NXP workstation to minimize blood layer disruption.

### Evaluating altering condition of qTRACE

Altering condition experiments were performed to characterize the tolerance of qTRACE to variations. For changing seeding density, D = 100 was used and the estimated seeding parasite in untreated wells ranged from 5-50. For changing D factor, an estimated 30 parasites per well in untreated wells were used.

### Dose response qTRACE

This method was designed to test the effect of compounds in a high throughput format using qTRACE. Compounds were tested in a 12-point semi log dilution in duplicates. Synchronized 0-3hpi parasites were seeded into 384 well plates with 50-100 parasites per well. The seeded plates were then drugged with the source plate using pintool affixed to the NXP (V&P Scientific). For single endpoint experiments, plates were fixed and stained at 120h. For time lapse experiments, whole plates were imaged using transmitted light at 48h interval followed by qTRACE AI analysis.

### Curve fitting

For experiments containing multiple timepoint, an exponential growth curve was fitted using the average colony area measured at each timepoint for each condition:

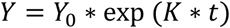

The fitted parameters are:

Y_0_: the best fitted starting value when t = 0.

K: the growth constant of the fitted curve for the condition and parasite line used

t: time

In the case of a comparison between two fitted curves, Y_0_ was restricted to untreated control or the reference parasite strain. K was extracted for calculation of GR.

In dose response experiments, SVI and GR of each concentration at each timepoint for each drug is calculated, then fitted into a sigmoidal curve:

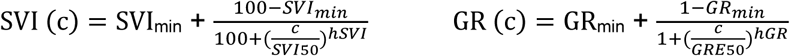

The fitted parameters are:

SVI_min_ or GR_min_: the maximum possible effect of a drug on parasite viability or growth inhibition in a fitted curve

hSVI or hGR: the hill slope coefficient of the fitted curve

SVI50 or GRE50: the concentration that confers half maximal effect on parasite viability or half maximal effect in GR curve.

### Imaging and quantification, and transmitted light imaging for deep learning

For fixed timepoints, plates were fixed and stained with 5µg/ml Hoechst and 1% glutaraldehyde. Imaging was performed using Lionheart FX (Biotek) or ImageXpress (Molecular device). Whole wells were captured for all 384 wells with 4X magnification. DAPI intensity and focus was determined using untreated wells as reference High content analysis was performed using image metrics including colony number, size, area, and signal intensity. We found a minimum size of 20 µm of colonies to allows for most optimal removal of small backgrounds and small artefacts. The number of colonies and the area of each colony were then exported for further analysis. Identified outliers were verified on the corresponding images and then discarded.

NF54-GFP parasite was used for the generation of deep learning training datasets. Whole well transmitted light images were acquired with 4X magnification using ImageXpress. The same sample was first imaged through transmitted light microcopy then immediately followed by GFP channel to acquire GFP reference image. In a continuous experiment, the plate was returned to culture condition and imaged again at next timepoint.

### Image processing and training data preparation

Original TIFF image with resolution of 2048×2048 and depth of 16-bit grayscale was first normalized to 2048×2048 resolution and 8-bit RGB format. The edge of every image was cropped by 150 pixels to remove areas with uneven illumination. All polygon annotations were initially performed by human professionals. Quality checks of all images and masks were performed with at least two rounds of random selections by two independent reviewers. To optimize computational efficiency and maintain consistent input dimensions across the dataset, all images and masks were downsampled to 512×512 pixels using the Lanczos resampling algorithm^32^. To enable rapid data loading during the iterative training process, all standardized images and their corresponding segmentation masks were subsequently converted to NumPy array format. The dataset was randomly partitioned into training (70%), validation (15%), and test (15%) sets.

### Deep Learning U-Net CNN model training

For semantic segmentation, we implemented a U-Net architecture comprising an encoder–decoder structure^33^. The encoder comprises four downsampling stages with 2×2 max pooling, progressively increasing the number of feature channels from 64 to 128, 256, and finally 512. Each stage uses two 3×3 convolutional layers with batch normalization and ReLU activation to capture contextual and morphological features^34–36^. At the bottleneck, feature maps are processed by a double convolution block with 1024 channels to capture high-level semantic features. The decoder mirrors the encoder with four upsampling stages using 2×2 transposed convolutions. Skip connections concatenate encoder features with decoder outputs to preserve spatial details, followed by double convolution blocks for refinement. A final 1×1 convolution produces the single-channel segmentation map for binary classification.

The model was trained using the Adam optimizer with an initial learning rate of 1×10⁻⁴ and an adaptive scheduler that reduced the rate after 5 stagnant validation epochs^37^. Training was conducted over 50 epoch with a batch size of 32, and validation performance was monitored to prevent overfitting. The final model was selected based on the best validation score, with all metrics reported on a held-out test set. Visual explanations of the model is generated using Grad-CAM to derive attention heatmaps across model layers^38^.

### Colony quantification

Following segmentation, automated quantification of colony number and area was performed on the predicted binary masks, where white regions (pixel value = 1) represent identified colonies and black regions (pixel value = 0) represent background. Colony enumeration was achieved through connected component labeling. We employed 8-connectivity (Moore neighborhood) for the connected component analysis, which considers pixels connected through both edges and corners as belonging to the same colony. The algorithm iteratively groups adjacent positive pixels until all connected regions are identified, with the total number of distinct labels corresponding to the colony count. Colony areas were quantified by calculating the sum of positive pixels within each segmented region. The total colonization density for each image was computed as the aggregate sum of all positive pixels in the binary mask.

### Performance evaluation and metrics

To evaluate colony enumeration, we used Mean Absolute Error (MAE), Root Mean Squared Error (RMSE), Exact Match Rate, tolerance-based accuracy (±1 and ±2 colonies), and the Pearson correlation coefficient to assess counting precision and consistency. For colony segmentation, we evaluated performance using standard pixel-wise metrics, including Intersection over Union (IoU), pixel accuracy, precision, recall, and F1 score, computed from binary masks of predicted and ground truth annotations.

### Vision transformer model for semantic segmentation

As an alternative approach to the U-Net architecture, we implemented a Vision Transformer (ViT) model adapted for semantic segmentation^39^. All 512×512 input images were divided into non-overlapping patches of 16×16 pixels, resulting in a sequence of 1024 patches (32×32 grid). To preserve spatial information, learnable position embeddings are added to the patch embeddings, which were then processed through a stack of transformer encoder layers, with layer normalization and residual connections.

For adapting the transformer to dense prediction, the sequence of output embeddings from the final transformer layer must be reshaped back to a 2D spatial representation and upsampled to match the original image resolution. This is achieved through a decoder head that first reshapes the patch tokens back to their spatial configuration, then applies successive upsampling layers to recover the full 512×512 resolution, ultimately producing pixel-wise binary predictions for colony segmentation. We followed the same training strategy and image processing settings for ViT model as well.

## Acknowledgements

We greatly appreciate Dr. David Fidock for the Dd2 K13 mutant lines and NF54 GFP line. We Gratefully acknowledge NIH (R21 AI168803) and Georgia Research Alliance for funding support. We thank Medicines for Malaria Venture for the reference anti-malarial compound library.

