## Supplementary Information for "High throughput quantitative tracking of *Plasmodium falciparum* clonal blood stage parasite growth and applications for antimalarial drug discovery"

**Supplementary Table 1. Summary of single-parasite viability index (SVI) data from qTRACE in *P. falciparum* dihydroartemisinin (DHA) exposure experiments.** Synchronous ring stage parasites were exposed to DHA (700 nM) for 6 hr, the drug was washed off, and the parasites were used to initiate qTRACE studies.

| | SVI day7 ± SEM | SVI day10 ± SEM | SVI day13 ± SEM | SVI day16 ± SEM | SVI day19 ± SEM | SVI day22 ± SEM |
|---|---|---|---|---|---|---|
| **Dd2** | 0.062±0.007 | 0.104±0.025 | 0.079±0.016 | 0.091±0.006 | 0.07±0.007 | 0.091±0.01 |
| **Dd2-K13-C580Y** | 4.323±0.851 | 4.958±0.543 | 5.255±1.158 | 5.438±0.792 | 5.494±0.573 | 4.918±0.422 |
| **Dd2-K13-R539T** | 8.189±1.066 | 9.937±1.334 | 11.115±2.549 | 11.055±1.249 | 8.889±1.081 | 9.425±1.204 |
| **D6** | 0.024±0.004 | 0.029±0.009 | 0.035±0.007 | 0.035±0.007 | 0.037±0.007 | 0.038±0.006 |
| **D6R** | 0.081±0.012 | 0.116±0.014 | 0.125±0.015 | 0.136±0.014 | 0.134±0.009 | 0.143±0.021 |
| **W2** | 0.237±0.01 | 0.354±0.039 | 0.428±0.044 | 0.47±0.06 | 0.501±0.085 | 0.504±0.081 |
| **W2R** | 0.536±0.037 | 0.794±0.138 | 0.808±0.101 | 0.798±0.113 | 0.841±0.136 | 0.803±0.08 |
| **C235** | 0.093±0.022 | 0.105±0.023 | 0.068±0.009 | 0.107±0.023 | 0.069±0.005 | 0.073±0.007 |
| **NHP-4781** | 0.18±0.043 | 0.224±0.037 | 0.264±0.09 | 0.253±0.042 | 0.273±0.05 | 0.251±0.052 |

**Supplementary Table 2. Summary of growth rate (GR) inhibition data from qTRACE in DHA exposure experiment.** K value is derived from the growth curve of each condition. In the case of fitness calculations, the non-DHA treated K value of each parasite were used to calculate GR value.

| | K ± SEM | K (DHA) ± SEM | GR ± SEM |
|---|---|---|---|
| **Dd2** | 0.15±0.004 | 0.13±0.003 | 0.81±0.019 |
| **Dd2-K13-C580Y** | 0.13±0.008 | 0.13±0.007 | 1.03±0.034 |
| **Dd2-K13-R539T** | 0.16±0.006 | 0.16±0.003 | 1.038±0.043 |
| **D6** | 0.125±0.003 | 0.11±0.002 | 0.85±0.01 |
| **D6R** | 0.124±0.002 | 0.127±0.002 | 1.03±0.002 |
| **W2** | 0.15±0.011 | 0.12±0.007 | 0.79±0.028 |
| **W2R** | 0.12±0.007 | 0.12±0.006 | 1.01±0.014 |
| **C235** | 0.15±0.004 | 0.13±0.002 | 0.88±0.013 |
| **NHP-4781** | 0.1±0.006 | 0.09±0.004 | 0.85±0.048 |
| **Dd2-K13-C580Y vs Dd2 fitness** | – | – | 0.73±0.042 |
| **Dd2-K13-R539T vs Dd2 fitness** | – | – | 0.975±0.027 |
| **D6R vs D6 fitness** | – | – | 0.87±0.049 |
| **W2R vs W2 fitness** | – | – | 0.7±0.019 |

**Supplementary Table 3. Summary of 50% single-parasite viability index (SVI50) from qTRACE experiments with Dd2 and 15 antimalarial drugs; 50% inhibitory concentrations (IC50) from bulk measurements with each drug are shown for comparison. (Units = nM).**

| | Artesunate | Dihydroartemisinin | Atovaquone | Artemether | OZ439 |
|---|---|---|---|---|---|
| **SVI50±SEM** | 0.26±0.01 | 0.46±0.10 | 1.26±0.56 | 2.27±0.55 | 1.17±0.27 |
| **IC50±SEM** | 0.26±0.001 | 0.3±0.02 | 3.02±0.73 | 2.31±0.18 | 1.97±0.288 |
| | **Amodiaquine** | **Lumefantrine** | **Mefloquine** | **DSM265** | **Azithromycin** |
| **SVI50±SEM** | 0.25±0.022 | 2.78±1.13 | 2.69±0.91 | 46.22±15.93 | 242.457±69.39 |
| **IC50±SEM** | 0.44±0.012 | 4.85±0.65 | 6.29±1.65 | 48.35±3.31 | 81.42±21.47 |
| | **Pyronaridine** | **MMV390048** | **KAE609** | **Pyrimethamine** | **Sulfadoxine** |
| **SVI50±SEM** | 1.12±0.30 | 28.3±14.23 | 0.68±0.11 | >10 | >50 |
| **IC50±SEM** | 1.72±0.04 | 22±3.53 | 0.81±0.05 | >10 | >50 |

**Supplementary Table 4. Summary of deep learning model performance of U-Net and Vision Transformer architectures (ViT) for both colony counting and area measurements.**

| Quantification | Metrics* | U-Net | ViT |
|---|---|---|---|
| **Count** | **MAE** | 2.63 | 2.87 |
| | **RMSE** | 4.8 | 5.66 |
| | **Exact Match** | 24.19% | 20.67% |
| | **Within ±1 Count** | 51.16% | 46.38% |
| | **Within ±2 Count** | 67.36% | 55.93% |
| | **Correlation** | 0.9541 | 0.9347 |
| **Area** | **IoU** | 0.7445 | 0.6988 |
| | **Pixel Accuracy** | 0.9983 | 0.9956 |
| | **Precision** | 0.8611 | 0.8053 |
| | **Recall** | 0.8461 | 0.7890 |
| | **F1 score** | 0.8535 | 0.7971 |

*Mean absolute error = MAE; Root mean square error = RMSE; Intersection over union = IoU.

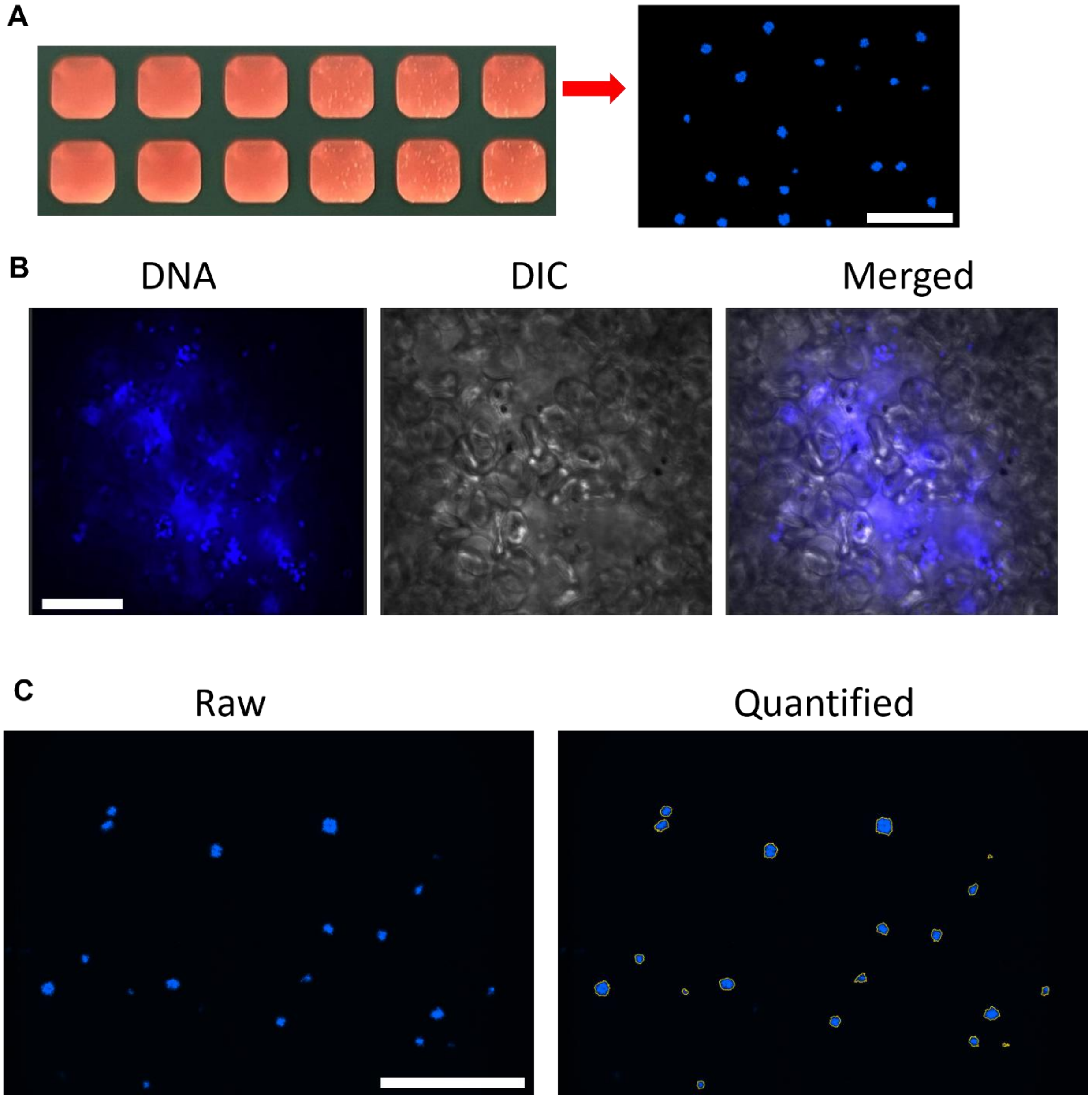


**Fig. S1. Representative examples of parasite colonies and how they were quantified. (A)** Representative colony observation results and well view from an experiment. Scale bar 1000μm. **(B)** 100X magnification images of the same colony shown in Supplementary Video 2. Scale bar 15μm. **(C)** Representative quantification of colonies in one well in 384 well plates. Scale bar 1000μm.

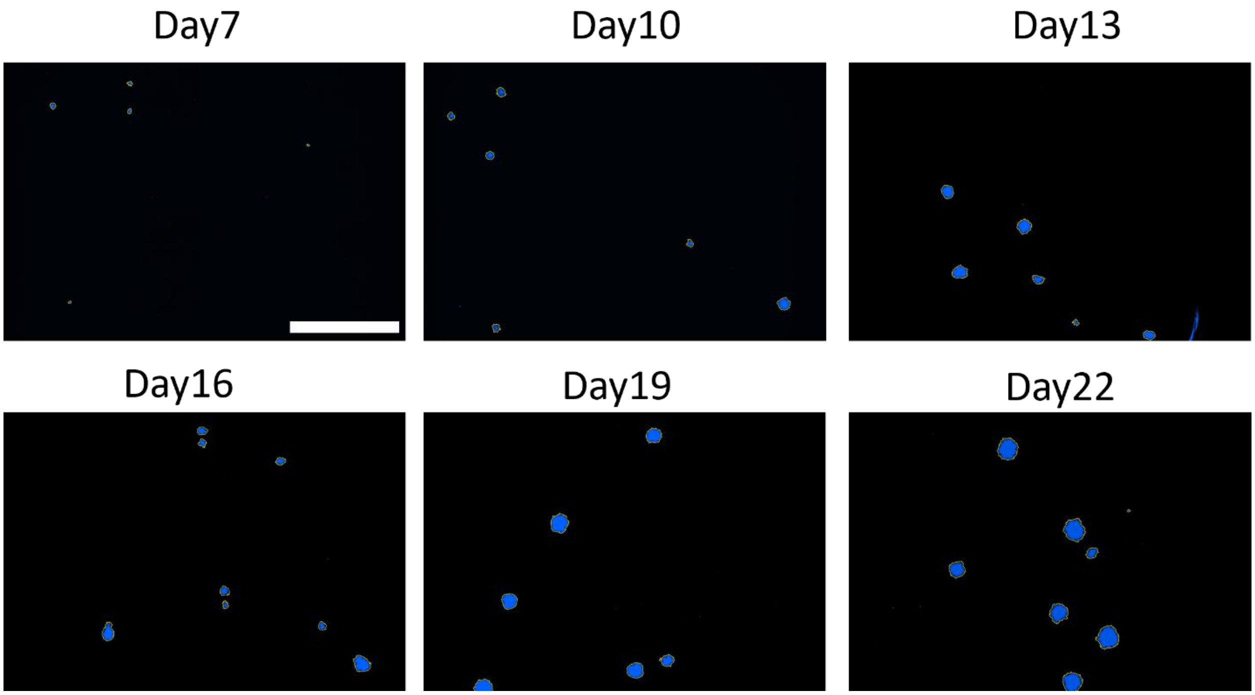


**Fig. S2. Quantification of colony numbers and sizes at different timepoints with DNA dye (Hoechst 33542).** Quantification of colonies and growth measurements of each colony. Most of the fluorescent artifacts can be filtered such as ones shown on day 10 and day 13. Scale bar 1000μm.

**A**

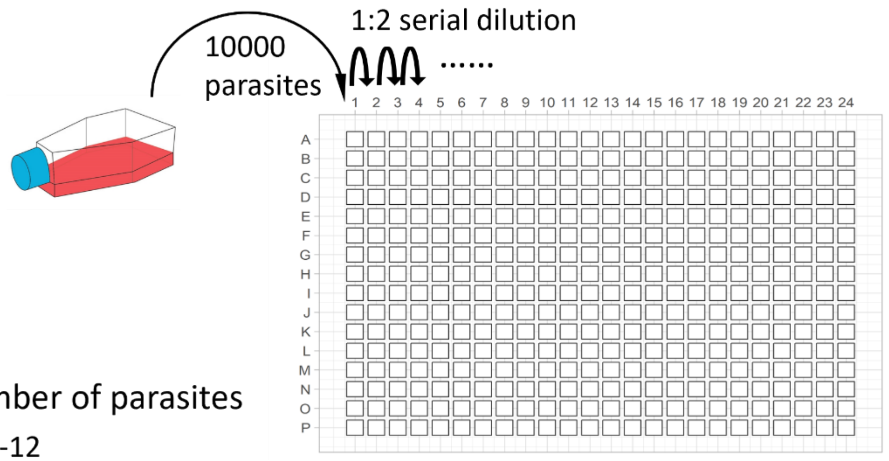


Estimated number of parasites

Dilution point 1-12

| 1 | 2 | 3 | 4 | 5 | 6 | 7 | 8 | 9 | 10 | 11 | 12 |
|---|---|---|---|---|---|---|---|---|---|---|---|
| 10,000 | 5000 | 2500 | 1250 | 625 | 312.5 | 156.25 | 78.125 | 39.063 | 19.531 | 9.766 | 4.883 |

Dilution point 13-24

| 13 | 14 | 15 | 16 | 17 | 18 | 19 | 20 | 21 | 22 | 23 | 24 |
|---|---|---|---|---|---|---|---|---|---|---|---|
| 2.441 | 1.221 | 0.610 | 0.305 | 0.153 | 0.076 | 0.038 | 0.019 | 0.010 | 0.005 | 0.002 | 0.001 |

**B**

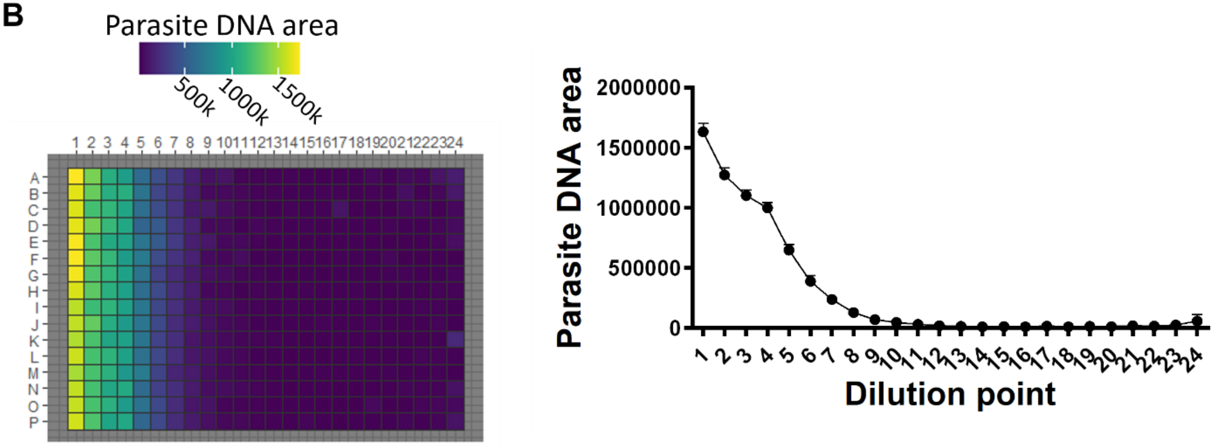


**Fig. S3. Validation of colony quantification using serial dilution of parasites in a 384 well plate. (A)** Illustration of the two-fold serial dilution experiment and estimated number of parasites per well in each column. **(B)** Total DNA content in each well (left) and average DNA content in every dilution point (right).

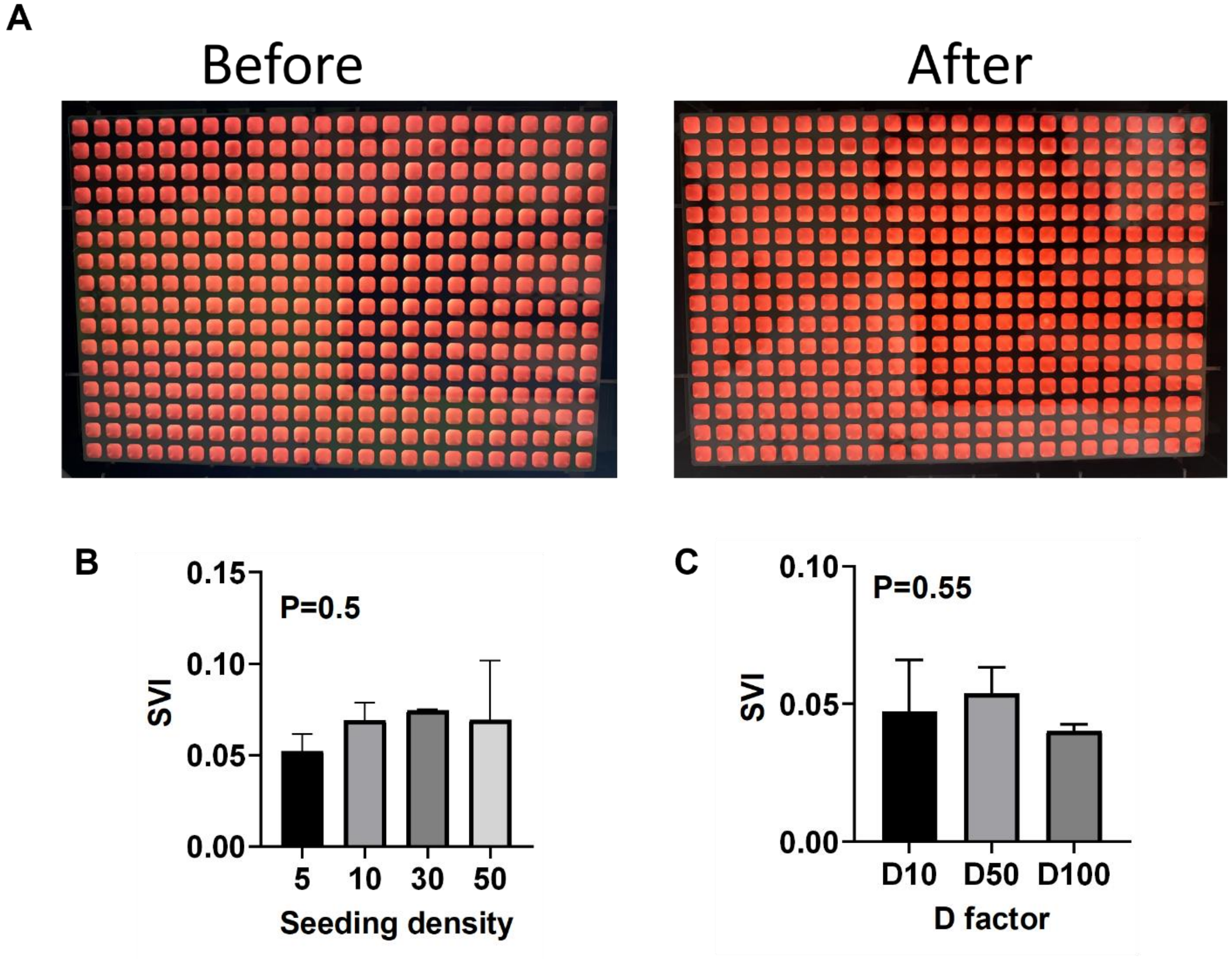


**Fig. S4. qTRACE protocol tolerance. (A)** No visual disruption on the static blood layer after media change through designed robotic protocol. This is a critical step to ensure the colony count accurately reflects original starting parasites at seeding. **(B)** Effect of altering cell density with D=100 on single-parasite viability index (SVI) assessments after DHA exposure. P-value calculated from two-way Anova. **(C)** Effect of altering dilution (D) factors on SVI assessments after DHA exposure. P-value calculated from two-way Anova.

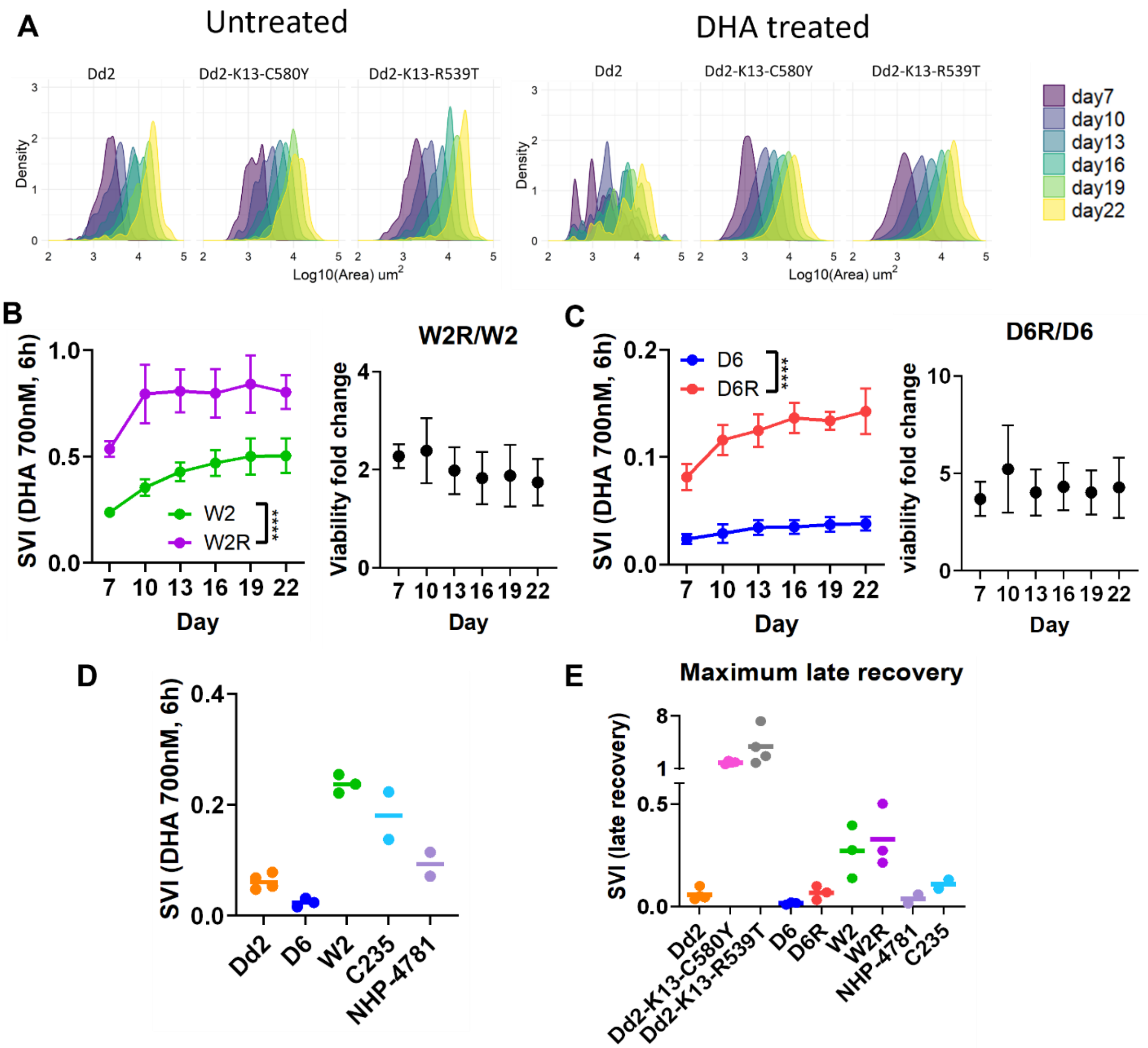


**Fig. S5. qTRACE reveals DHA susceptibility and delayed recovery post day 7 from artemisinin induced dormancy. (A)** Colony growth of Dd2, Dd2-K13-C580Y and Dd2-K13-R539T from day 7 to 22. Data are from two biological replicates. **(B)** Time-dependent single-parasite viability index (SVI) of W2 and W2R at continuous timepoints (left) and fold change in SVI between W2 and W2R across different timepoints (right). n=80 Technical Replicate and n=3 Biological Replicate. P-value calculated from two-way Anova. **(C)** Time-dependent SVI of D6 and D6R at continuous timepoints (left) and fold change in SVI between D6 and D6R across different timepoints (right). n=80 Technical Replicate and n=3 Biological Replicate. P-value calculated from two-way Anova. **(D)** Survival SVI assessments of five artemisinin sensitive parasites on day 7. n=80 Technical Replicate and n=3 Biological Replicate. **(E)** Maximum late recovery SVI (Max SVI-Survival SVI) of nine tested parasites. More details in Methods. Additional data is shown in Supplementary Table 1.

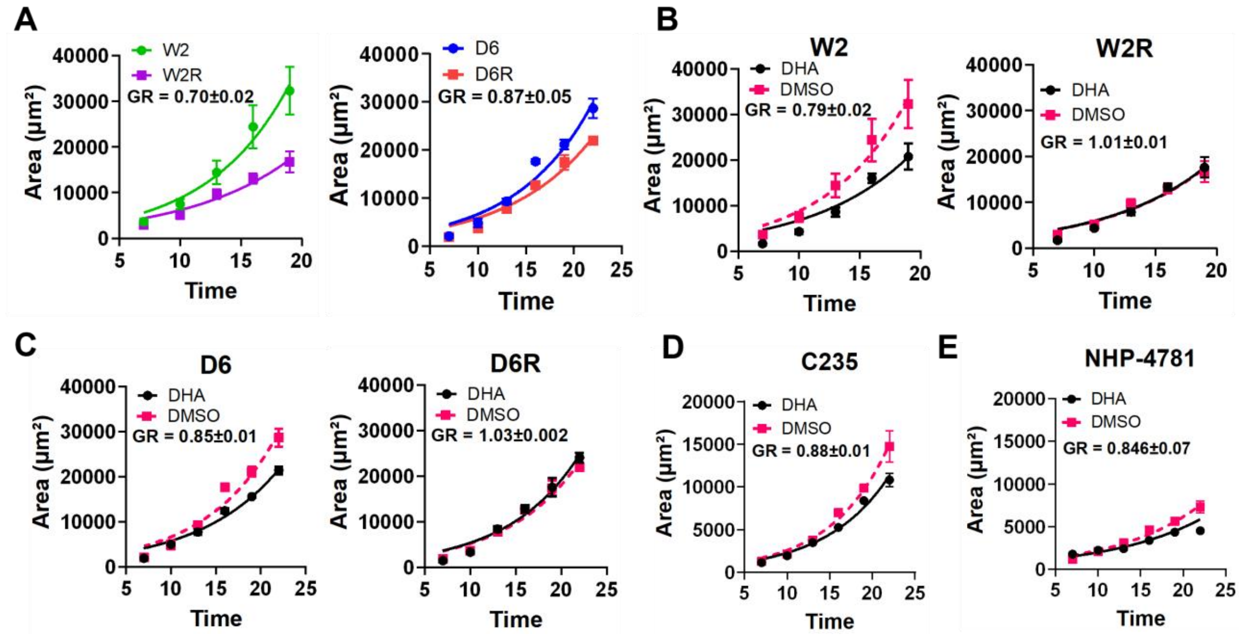


**Fig. S6. qTRACE reveals growth inhibition imposed by drug exposure and fitness differences between parasite lines. (A)** Fitted growth curves of averaged colony area at different timepoints and calculated growth rate (GR) without any treatment for W2 and W2R (left) or for D6 and D6R (right). (n=80 technical replicates and n=3 biological replicates) **(B)** Fitted growth curves of averaged colony area at all timepoints in control (DMSO) and after exposure to DHA; GR calculated by comparing the two conditions for W2 (left) and W2R (right). **(C)** Identical studies (as shown in C were conducted with D6 (left) and D6R (right), **(D)** C235 and **(E)** for NHP-4781. Data in C-F included n=80 technical replicates and n=3 biological replicates for each *P. falciparum* strain. Data for additional parasites is shown in Supplementary Table 2. All P-values were calculated from two-way Anova.

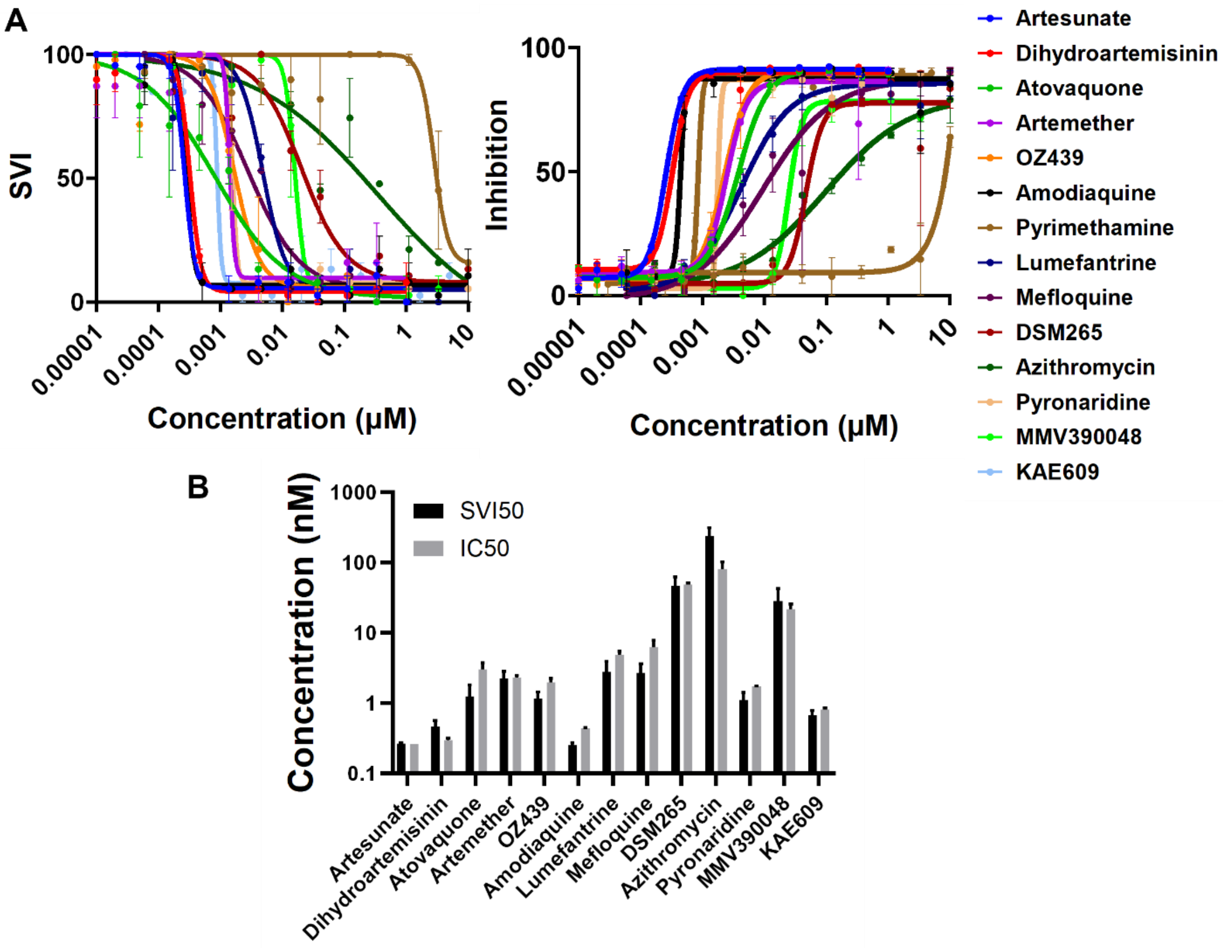


**Fig. S7. Application of qTRACE in drug testing. (A)** Dose response curves from qTRACE (left) and conventional bulk methods (right) for 14 drugs with a 12-point semi log concentration at 120h. (Error bar=SEM and n=2 biological replicates); **(B)** 50% single-parasite viability index (SVI50) and 50% inhibitory concentrations (IC50) for 13 drugs. Pyrimethamine was excluded. (Error bar=SEM and n=2 biological replicates). More data in Supplementary Table 3.

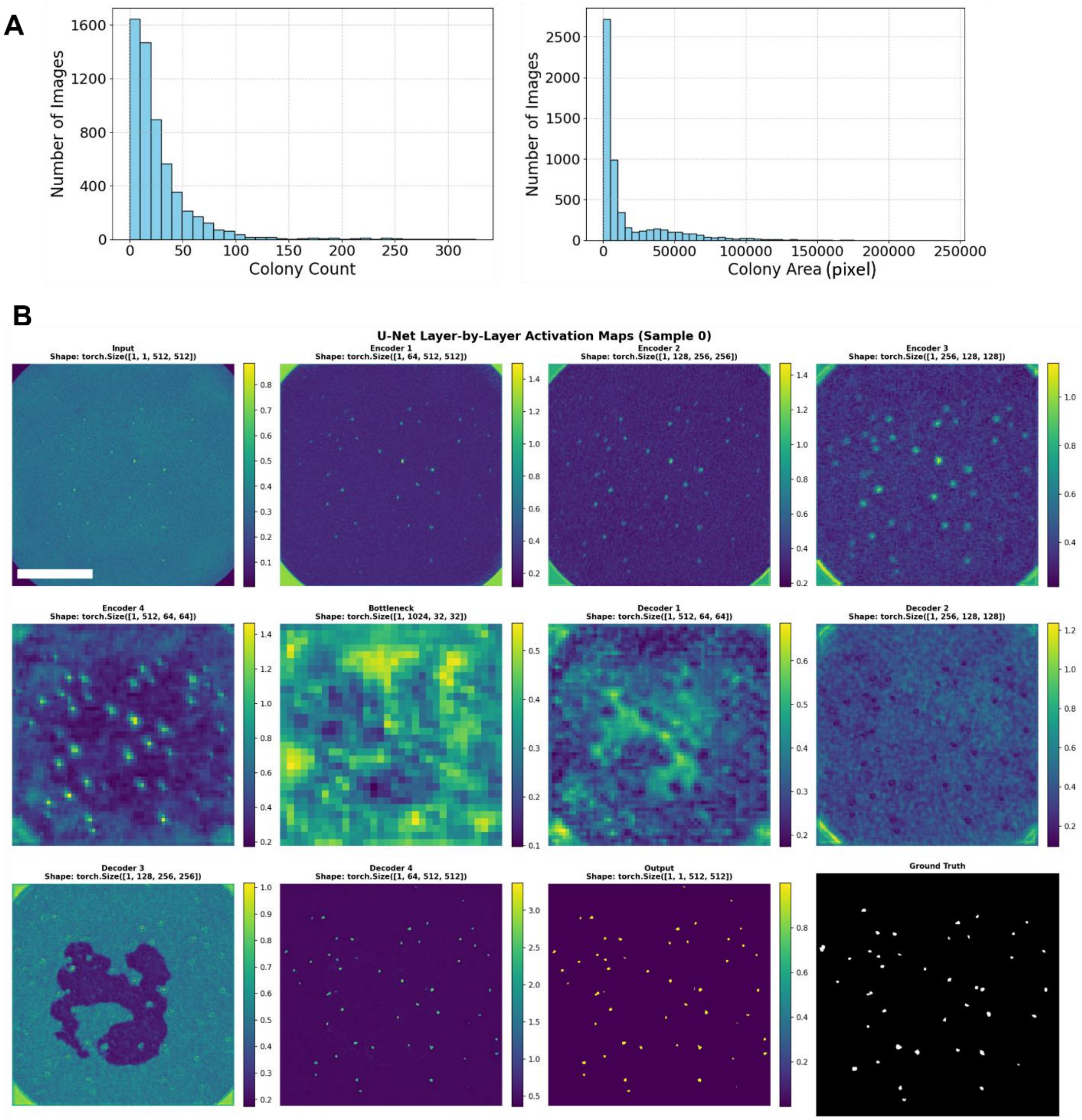


**Fig. S8. Deep learning U-Net convolutional neural network model predicts colony quantification. (A)** Distribution of colony count (left) and colony area (right) of training datasets. **(B)** Attention map of a sample image in each layer throughout qTRACE AI model U-Net CNN model. Heatmaps show feature activations progressing from input through encoder (1-4), bottleneck, and decoder (1-4) layers to output, with color intensity indicating activation magnitude. Tensor dimensions (batch, channels, height, width) are noted above each panel. Ground truth segmentation is shown for reference (bottom right).

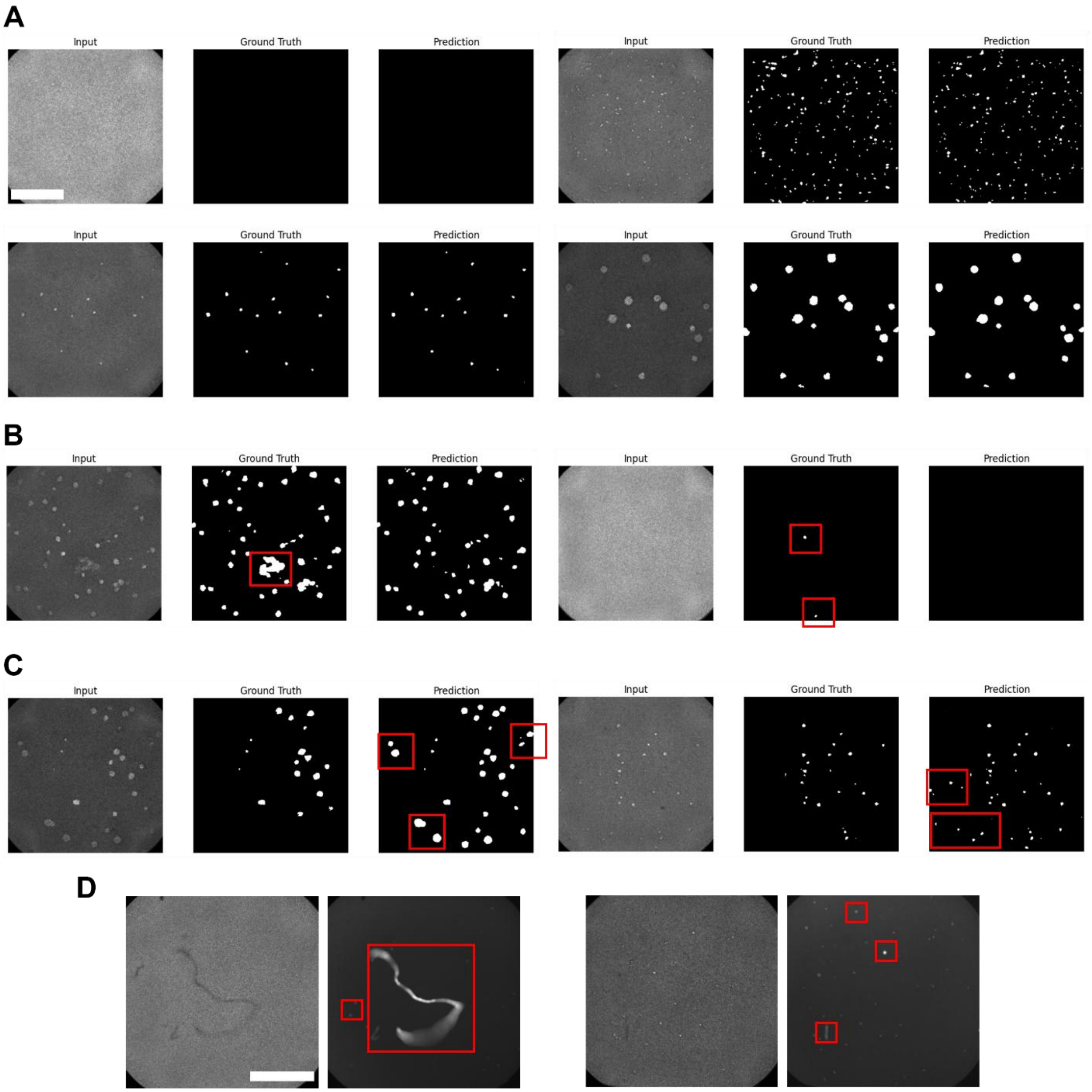


**Fig. S9. Representative image sets demonstrating qTRACE AI mode performance. (A)** Images showing highly identical analysis from annotated results and model predicted results across diverse conditions, including empty (top left), high seeding density (top right), low seeding density (bottom left) and later timepoint (bottom right). **(B)** Images showing correction of false positives and debris by trained model. Red box indicating false positives from human annotation but corrected in model prediction. **(C)** Images showing correction of false negatives by trained model. Red box indicating false negatives missed from human annotation but corrected in model prediction. **(D)** Images showing noises in GFP fluorescent images and corresponding transmitted light images. Red box indicates artifacts in GFP images that do not appear in transmitted light images. Scale bar = 1000μm for all images.

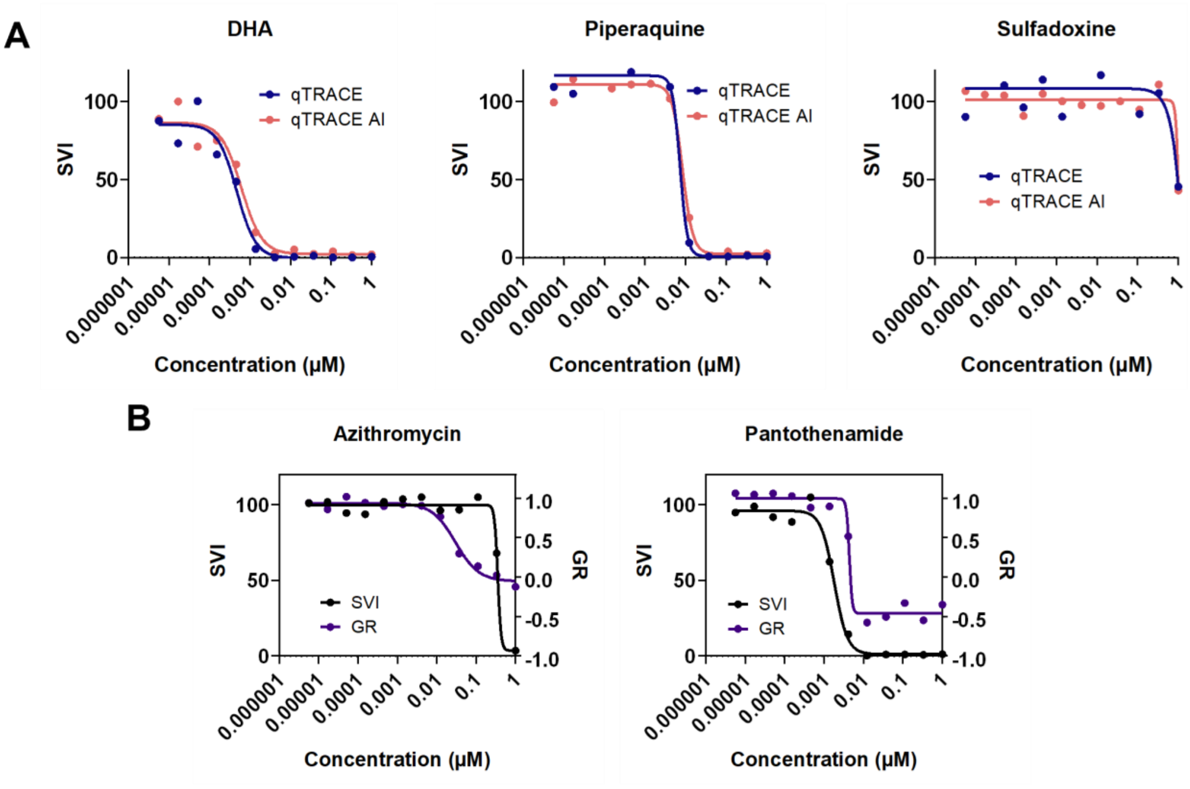


**Fig. S10. qTRACE analysis reveals cytostatic and cytotoxic drug effects. (A)** Dose response curve of 3 drugs showing similar results from qTRACE AI and DNA stain qTRACE. **(B)** Dose response curve of two drugs showing strong cytostatic effect (left) and cytotoxic effect (right).
